# iTRAQ-based proteomic and phosphoproteomic analyses of STRIPAK mutants from the fungus *Sordaria macrospora* identifies a conserved serine phosphorylation site in PAK kinase CLA4 to be important for sexual development and polarized growth

**DOI:** 10.1101/828111

**Authors:** R Märker, B Blank-Landeshammer, A Beier-Rosberger, A Sickmann, U Kück

**Affiliations:** Lehrstuhl für Allgemeine und Molekulare Botanik, Ruhr-Universität, 44780 Bochum, Germany; Leibniz-Institut für Analytische Wissenschaften-ISAS-e.V., Otto-Hahn-Straße 6b, 44227 Dortmund, Germany

**Keywords:** STRIPAK, iTRAQ-based proteomic and phosphoproteomic analyses, CLA4, a p21-activated kinase, *Sordaria macrospora*, sexual development

## Abstract

The highly conserved striatin-interacting phosphatases and kinases (STRIPAK) complex regulates phosphorylation of developmental proteins in eukaryotic microorganisms, animals, and humans. To first identify potential targets of STRIPAK, we performed extensive isobaric tags for relative and absolute quantification (iTRAQ)-based proteomic and phosphoproteomic analyses in the filamentous fungus *Sordaria macrospora.* In total, we identified 4,193 proteins and 2,489 phosphoproteins, which are represented by 10,635 phosphopeptides. By comparing phosphorylation data from wild-type and mutants, we identified 228 phosphoproteins to be regulated in all three STRIPAK mutants, thus representing potential targets of STRIPAK. To provide an exemplarily functional analysis of a STRIPAK-dependent phosphorylated protein, we selected CLA4, a member of the conserved p21-activated kinase (PAK) family. Functional characterization of the Δcla4 deletion strain showed that CLA4 controls sexual development and polarized growth. To determine the functional relevance of CLA4 phosphorylation and the impact of specific phosphorylation sites on development, we next generated phospho-mimetic and -deficient variants of CLA4. This analysis identified (de)phosphorylation of a highly conserved serine (S685) residue in the catalytic domain of CLA4 as being important for fungal cellular development. Collectively, these analyses significantly contribute to the understanding of the mechanistic function of STRIPAK as a phosphatase and kinase signaling complex.

## Introduction

The striatin-interacting phosphatases and kinases (STRIPAK) complex is a highly conserved macromolecular signaling complex that is involved in numerous cellular and developmental eukaryotic processes. In humans, malfunctions of STRIPAK subunits are linked with various diseases and cancer; whereas, in eukaryotic microorganisms, several processes, such as cell fusion, fruiting body formation, as well as symbiotic and pathogenic interactions, are controlled by STRIPAK (Kück et al. 2019). Although architecture, function, and regulation of STRIPAK are well characterized in diverse experimental systems (Hwang and Pallas 2014; Kück et al. 2016; Shi et al. 2016), our understanding of how STRIPAK regulates the phosphorylation of developmental proteins is still rudimentary. Further, it also currently remains elusive as to how the inactivation or inhibition of subunits of the STRIPAK complex affect the phosphorylation status of STRIPAK target proteins. Previously, we have used a large set of mutant and deletion strains to identify and structurally and functionally characterize STRIPAK subunits in the model fungus *Sordaria macrospora.* In extensive tandem affinity purification followed by mass spectrometry (TAP-MS) and yeast-two-hybrid analyses, we identified the structural (PP2AA) and catalytic (PP2Ac) subunits of PP2A, the B‴ regulatory subunit of PP2A (striatin), the striatin-interacting protein PRO22, SmMOB3, which is homologous to the mammalian vesicular trafficking protein Mob3, and PRO45, a homologue of the mammalian sarcolemmal membrane-associated protein SLMAP (Kück et al. 2016; Kück et al. 2019). Recently, a STRIPAK complex interactor 1 (SCI1) was identified that exhibited structural similarity to SIKE, a coiled-coil protein that serves as a negative regulator of pathological cardiac hypertrophy in humans (Reschka et al. 2018). Further, two STRIPAK-associated germinal center kinases (GCKIII) were also functionally characterized (Frey et al. 2015; Radchenko et al. 2018), linking STRIPAK with the fungal septation initiation network (SIN). Of interest is that SIN is homologous to the animal HIPPO pathway, which is negatively regulated by STRIPAK.

Quantitative MS-based phosphoproteomics offers the possibility to monitor thousands of phosphorylation sites even from minute sample amounts. This approach enables the large-scale screening and identification of potential targets of phosphorylation or dephosphorylation by identification of the modified peptides with high confidence (Pagel et al. 2015). Besides label-free quantification and metabolic labelling (e.g. SILAC), reporter-ion-based quantification by usage of isobaric chemical labels (e.g. iTRAQ, TMT) has emerged as an effective technique for large-scale relative protein quantification. Due to the fact that iTRAQ allows simultaneous analysis of up to eight protein samples from different experimental conditions, this technique minimizes random errors during sample preparation, and therefore, proved to be especially suitable for phosphoproteomic analyses (Sachon et al. 2006; Wiese et al. 2007; Yang et al. 2007).

Here, we performed extensive iTRAQ-based proteomic and phosphoproteomic analyses to identify differentially (de)phosphorylated proteins, by comparing the wild-type phosphoproteome with those from three mutants lacking distinct subunits of STRIPAK. Phosphorylation sites in 228 proteins were differentially regulated in all three STRIPAK mutants. Among these, we identified CLA4, a member of the p21-activated kinase (PAK) family. Our analyses revealed a differentially phosphorylated residue, which is highly conserved in PAKs from human to eukaryotic microorganisms. The functional relevance of these phosphorylation sites on fungal development was tested using both phospho-mimetic and -deficient variants of CLA4. Our study provides an in-depth quantitative phosphoproteomics data set and extends the number of phosphorylated proteins that are putative targets of STRIPAK. Finally, we present the first evidence that STRIPAK regulates phosphorylation of a kinase from the PAK family.

## Results

In this work, detailed iTRAQ-based proteomic and phosphoproteomic analyses of the wild-type and three different STRIPAK deletion strains led to the identification of putative dephosphorylation targets of the STRIPAK complex in *S. macrospora*. Importantly, functional characterization of the PAK kinase CLA4, a target protein of STRIPAK deciphers their role in fungal cellular development.

### iTRAQ-based proteomic and phosphoproteomic analyses revealed putative dephosphorylation targets of the STRIPAK complex

To identify dephosphorylation targets of STRIPAK, we performed iTRAQ-based LC-MS/MS proteomic and phosphoproteomic analyses to directly compare relative changes in protein expression (Fig. 1). The iTRAQ strategy is based on the N-terminal labelling of peptides with up to eight amino group-reactive reagents coupled to different reporter and balancer groups (Ross et al. 2004; Wu et al. 2006). Thus, we were able to conduct a simultaneous analysis of up to eight different cultural samples. While these isobaric tags have the same mass, fragmentation during tandem MS (MS/MS) leads to the generation of mass-specific reporter ions whose abundances are used for protein quantification. Using the same sample for the simultanous analysis of both the global protein expression as well as the phosphoproteome further allows a better differentiation of true changes in phosphorylation levels from mere fluctuations of the associated proteins (Solari et al. 2016). For proteomic and phosphoproteomic analyses, we used probes from wild-type and three different STRIPAK deletion mutants that lacked either *pro11*, *pro22*, or *pp2Ac1* encoding the B‴ regulatory subunit of PP2A (PRO11), the catalytic subunit of PP2A (PP2Ac1), and the striatin-interacting protein PRO22, respectively (Bloemendal et al. 2012; Beier et al. 2016; Pöggeler and Kück 2006). All strains were grown under the same controlled growth conditions for three days in liquid surface cultures, containing bio malt maize media (BMM). As depicted in the workflow of Figure 1, we used two biological replicates per strain. After protein extraction and tryptic digestion, the peptides of each sample were labelled at the N-terminus with the different isobaric tags. After mixing the samples, high pH reversed-phase fractionation was performed for global proteome analysis, followed by nanoflow high-performance liquid chromatography (nano-HPLC) and MS/MS. In parallel, enrichment of phosphopeptides by titanium dioxide (TiO_2_) and fractionation by hydrophilic interaction liquid chromatography was conducted, followed by nano-HPLC and MS/MS. For both data sets, ratios of the reporter ion intensities of the deletion strains were calculated relative to the wild-type. To determine differentially regulated candidates, the total variation of ratios for each strain was determined and only phosphopeptides or proteins displaying a fold change of more than two times the standard deviation were considered as regulated.

**Figure 1:**
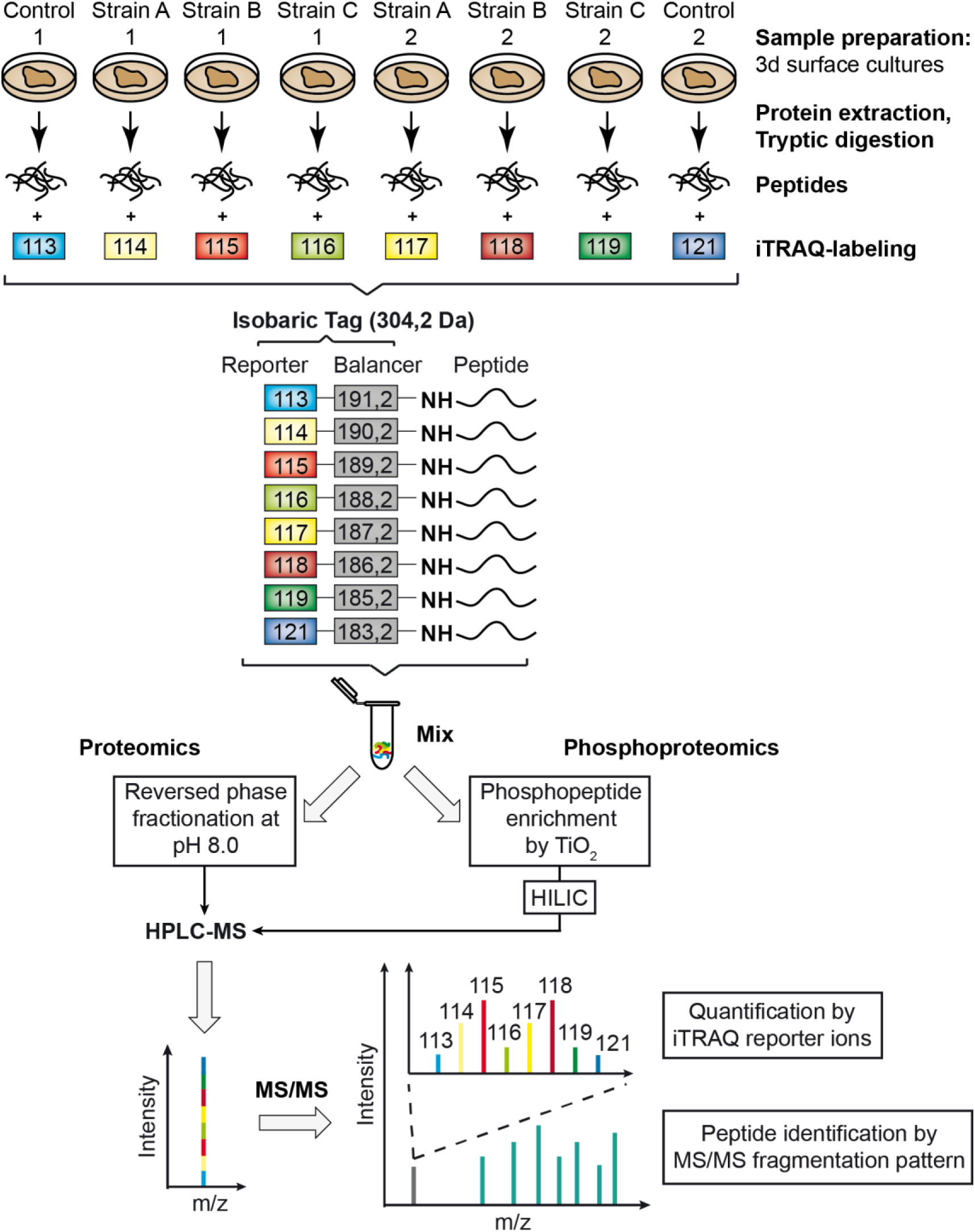
iTRAQ-based mass spectrometry workflow for proteomic and phospho-proteomic analysis of STRIPAK deletion mutants. For protein extraction, wild type and three deletion mutants were grown for three days. Two biological replicates were used for each strain. Following tryptic digestion, peptides of each sample were labeled at the N-terminus with isobaric tags for relative and absolute quantification (iTRAQ). After sample mixing, fractionation for the global proteome analysis and enrichment of phosphopeptides by titanium dioxide (TiO_2_) and hydrophilic interaction liquid chromatography (HILIC) were conducted. Subsequently, high-performance liquid chromatography (HPLC) and tandem mass spectrometry analysis (MS/MS) generated reporter ions with specific masses, which were used for quantification.

As shown in Figure 2, global proteome analysis of the wild-type and the deletion strains, namely, Δpro11, Δpro22, and Δpp2Ac1 revealed a total of 4,193 quantified proteins, identified with a minimum of two unique peptides per protein at 1% false discovery rate (FDR) on the peptide-spectrum match (PSM) level (Dataset S1). Phosphopeptide enrichment and fractionation led to the identification of 13,506 peptides, 11,161 of which harboured a single or multiple phosphorylations, indicating a relative enrichment efficiency of almost 85% on the PSM level. For 10,635 of those peptides, the phosphorylation site could be localized with high confidence (phosphoRS probability ≥ 90 %), cumulating in a total of 2,489 phosphoproteins (Dataset S2). Moreover, 39% of the phosphoproteins exhibited one phosphorylation site, while 61% contained multiple phosphorylation sites (Fig. S1A). Further, 79%, 20%, and 1% of the 10,635 peptides were phosphorylated on serine, threonine, and tyrosine residues, respectively (Fig. S1B). The proteomic and phosphoproteomic analyses showed an overlap of 1,727 proteins. Among them, 781 phosphoproteins were differentially phosphorylated or dephosphorylated at one or more phosphorylation sites. In addition, these proteins were stable in the global proteome, i.e. they showed similar abundances in the deletion strains and wild-type (Fig. 2).

**Figure 2:**
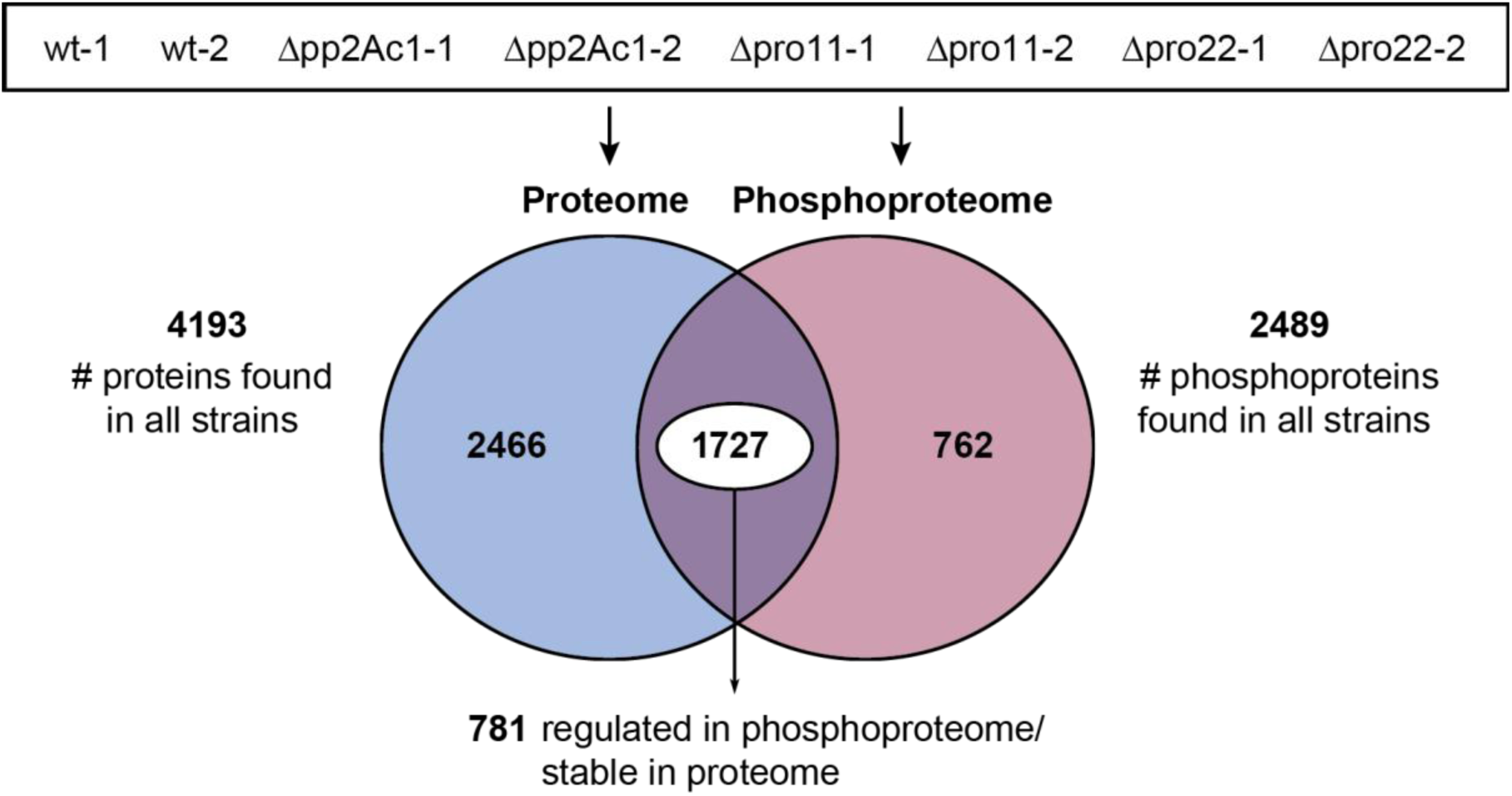
Proteins and phosphoproteins identified in the wild type and STRIPAK deletion strains. The total numbers of proteins and phosphoproteins from a proteomic and phosphoproteomic analysis of Δpro11, Δpro22, and Δpp2Ac1, as well as the wild type, are shown. The number in the intersection displays proteins detected in both analyses. Furthermore, the number of regulated phosphoproteins from all strains are given that were identified with similar abundances in the global proteome.

The GO annotation analysis of these phosphoproteins showed that numerous proteins play a role in metabolic processes and are located in organelles and the nucleus (Fig. S2). Thus, this study highlights that numerous proteins play a role in metabolic processes and are located in organelles and the nucleus (Fig. S2). Among the 781 phosphoproteins, 228 exhibited phosphorylation sites that were regulated in all three deletion strains (Fig. 3). The analysis of these putative STRIPAK targets revealed several proteins of known signal transduction pathways that exhibited up-or down-regulated phosphorylation sites (Dataset S3). The large number of identified phosphorylation sites permitted us to perform enrichment analyses for phosphorylation motifs of differentially regulated phosphorylation sites. Comparing the total proteome from *S. macrospora*, we found in up-regulated phosphosites six distinct motifs to be significantly enriched in all three knockout strains. The motifs [RXPpS] and [RXXpSP] showed the strongest enrichment (11.3- and 10.4-fold compared to the background; adjusted p-values 2.8E-06 and 1.5E-09, respectively) with the latter showing close resemblance to the reported substrate motif of kinases CDK5, ERK1/2, and GSK3 (Jaffe et al. 2004). The only enriched acidic motif [pSDXE] (5.2-fold enrichment, adj. p-value 1E-03) was reported to be a substrate of caseine kinase II (Schwartz and Gygi 2005). Interestingly, the fungi-specific phosphorylation motif [SXXpT] (Bai et al. 2017) was found to be neither significantly enriched among the up-regulated phosphorylation sites nor in the complete data set. A detailed overview of all enriched motifs can be found in the supplementary material (Data set S4).

**Figure 3:**
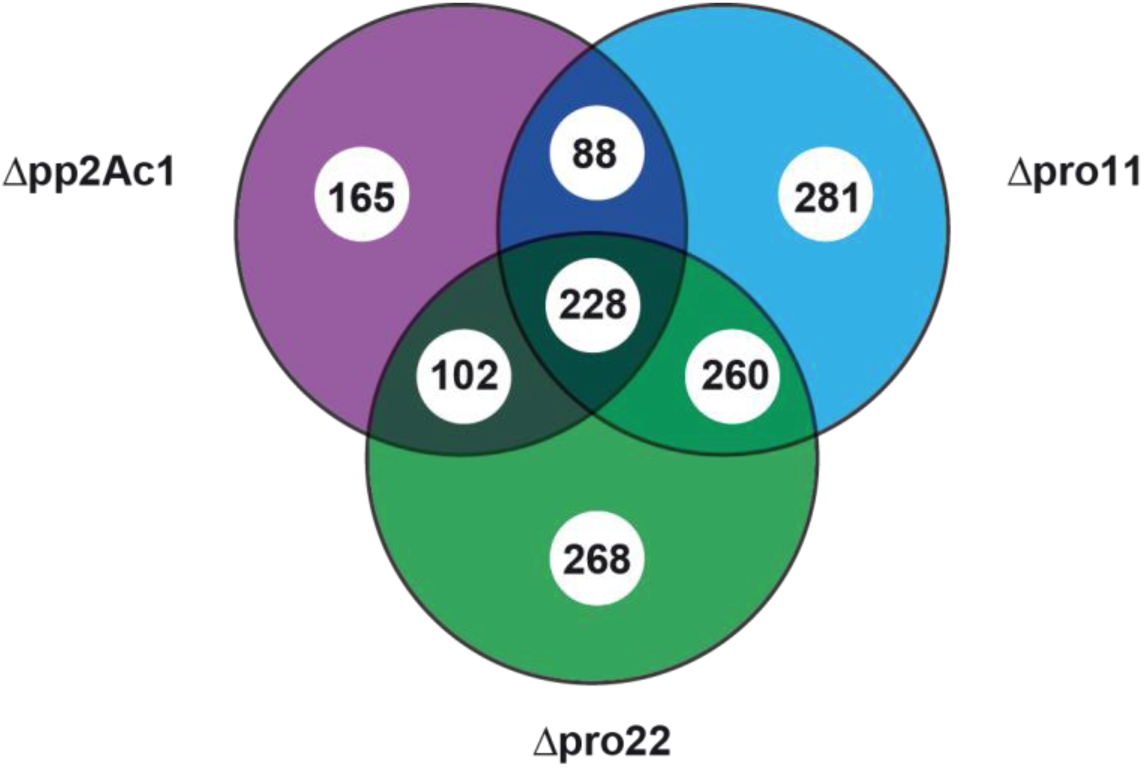
Venn diagram of 781 phosphoproteins with regulated phosphorylation sites in STRIPAK deletion strains. The comparative phosphoproteomic analysis of the wild type and the deletion strains Δpp2Ac1, Δpro11 and Δpro22 identified 781 phosphoproteins with regulated phosphorylation sites in the deletion strains. However, these proteins show similar abundances in all strains in the global proteome. The Venn diagram shows the number of phosphoproteins with regulated phosphorylation sites in each deletion strain. Numbers in intersections display phosphoproteins with phosphorylation sites that are regulated in two or three deletion strains. Some phosphoproteins are given in more than one intersection due to the fact that they exhibit multiple regulated phosphorylation sites.

### The PAK kinase CLA4 is a potential target of STRIPAK that controls sexual development and polarized growth

In Table 1, we present 29 selected proteins belonging to the functional classes of sexual signalling, kinases, phosphatases, and transcription factors. Among these are pheromone-processing peptidases KEX1 and KEX2, and the developmental protein HAM9, a target of transcription factor PRO1, which is involved in fruiting body formation and cell fusion (Mayrhofer and Pöggeler 2005; Fu et al. 2011). Further, we identified several transcription factors some of which were described to regulate fungal sexual development. Included in the list of predicted STRIPAK targets are phosphatases and kinases, which were described in several cases to be involved in fungal development. This is also true for CLA4, a member of the p21-activated kinases, which is conserved from humans to eukaryotic microorganisms. This class of kinases was also characterized for several filamentous fungi and is involved in a large variety of cellular processes (Boyce and Andrianopoulos 2011). However, an association with STRIPAK has not been described for any model organism, and functional analysis of phosphorylation sites in filamentous fungi is currently missing. Therefore, we provided a detailed functional analysis of CLA4, a potential target of STRIPAK.

**Table 1:**
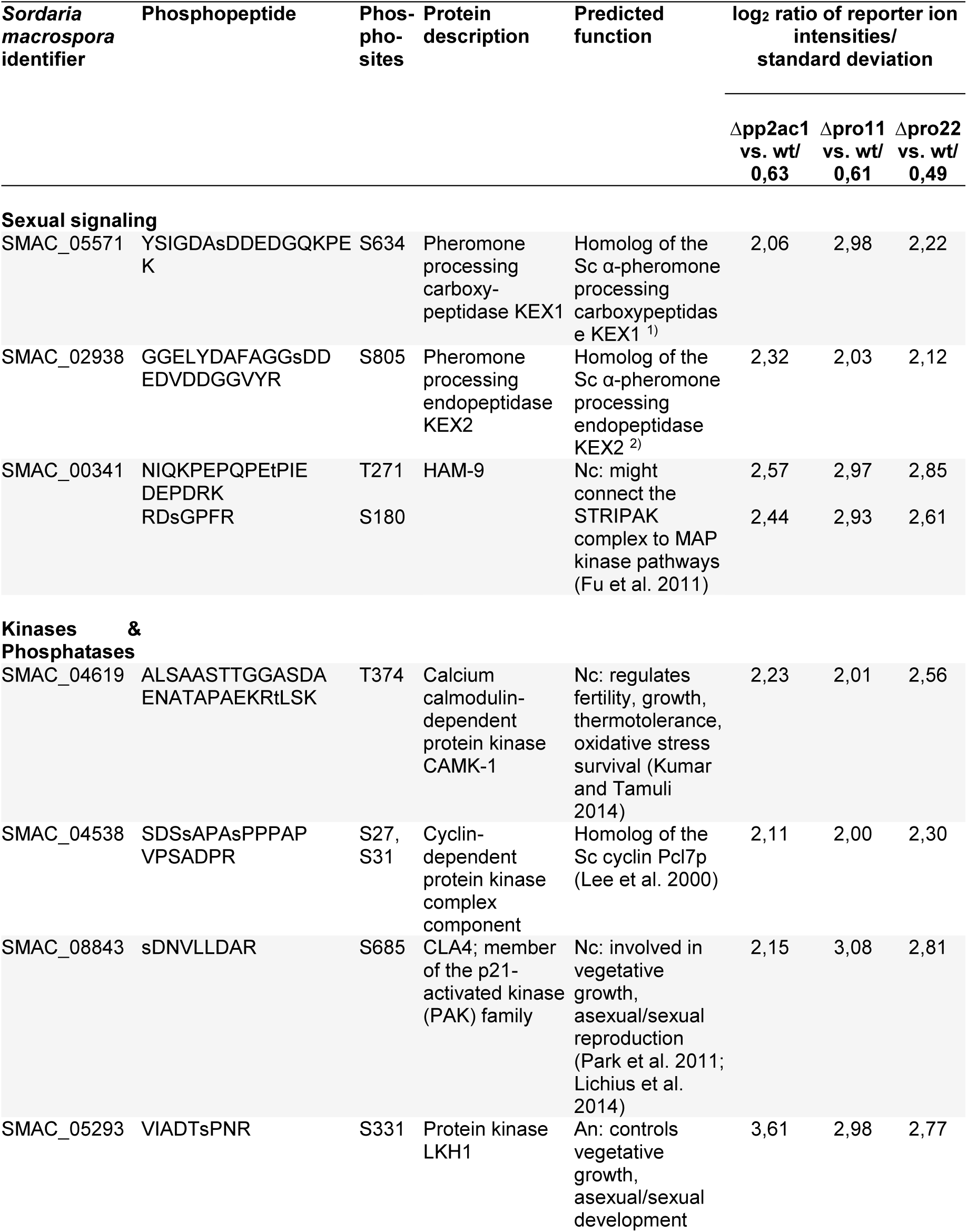
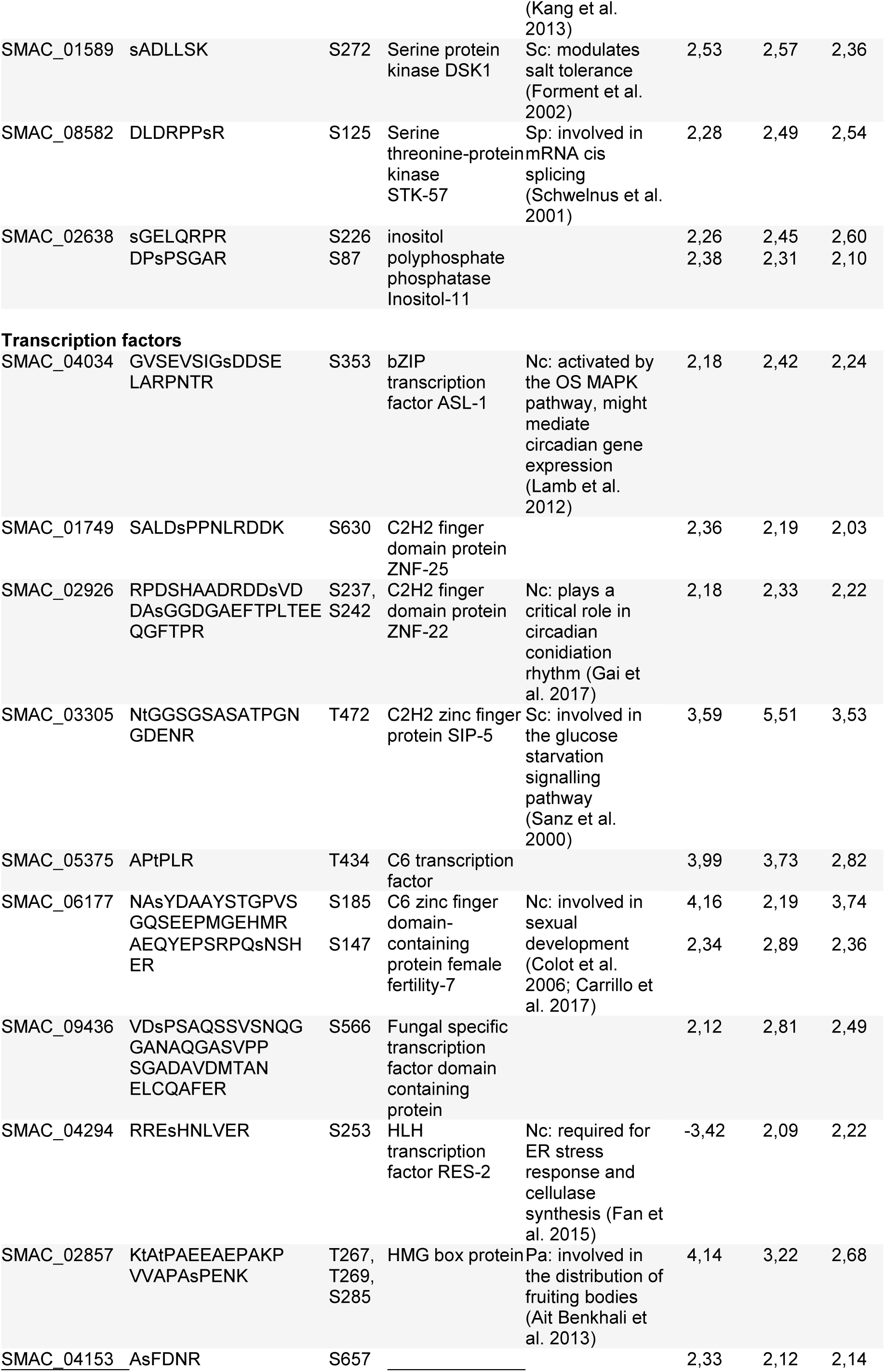
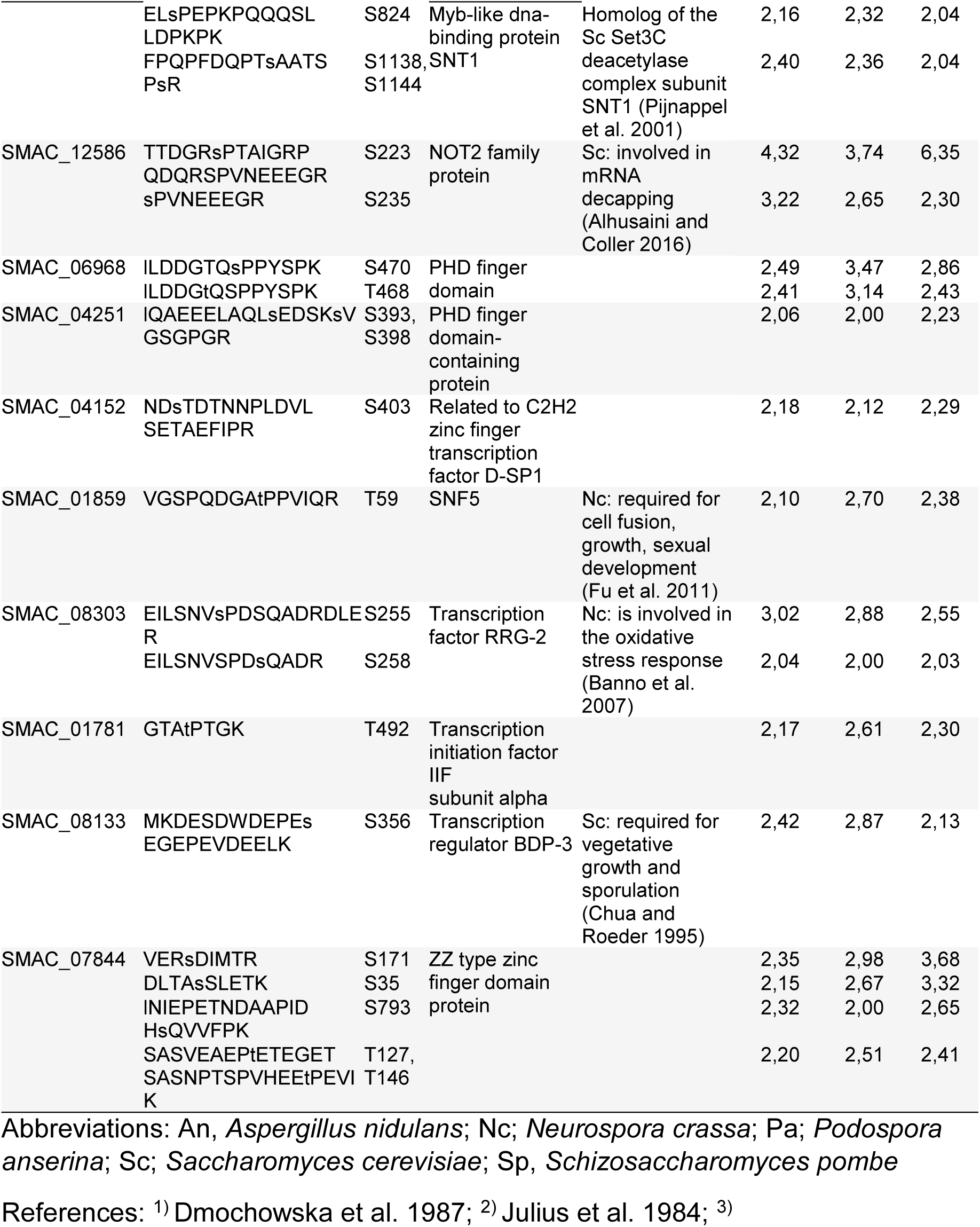
Phosphoproteins that appear to be regulated in all three investigated STRIPAK mutants Δpp2Ac1, Δpro11 and Δpro22. Given are phosphorylated peptides from a total of 29 selected proteins. For each phosphorylation site, log_2_ ratio of reporter ion intensity in deletion strain and wild type relative to the respective standard deviation are given.

The *S. macrospora SMAC_08843* gene comprises an open reading frame (ORF) of 2817 bp and is located at scaffold 236 of the *S. macrospora* genome v03 (Blank-Landeshammer et al 2019). The predicted protein has a length of 834 amino acids and shows high amino acid sequence identity to homologous CLA4 proteins in *Neurospora crassa* (EAA28056.1, 96.65% identity), *Fusarium graminaerum* (SCB65154.1, 69.92% identity), and *Magnaporthe grisea* (AAL15449.2, 69.36% identity) (Fig. S3). The protein exhibits 53.81% identity to *Saccharomyces cerevisiae* Cla4p (CAA96216.1) and 58.82% identity to human PAK1 (AAC50590.1). The conserved domain search using the Conserved Domain Search service (CD Search) at NCBI revealed a pleckstrin homology domain (PH), a p21-binding domain (PBD), and a kinase domain. These domains were present in the *S. cerevisiae* homologue Cla4p (Fig. 4A). The alignment of the amino acid sequences from *S. macrospora* and other filamentous fungi indicated that among the three identified phosphorylation sites in *S. macrospora*, only S685 is located in a highly conserved region within the kinase domain (Fig. 4B). Importantly, this phosphorylation site has not been functionally characterized in any organism to date, while the related amino acid is located within a substrate-binding site, and thus, seems to be of relevance (Fig. 4B, Fig. S3).

**Figure 4:**
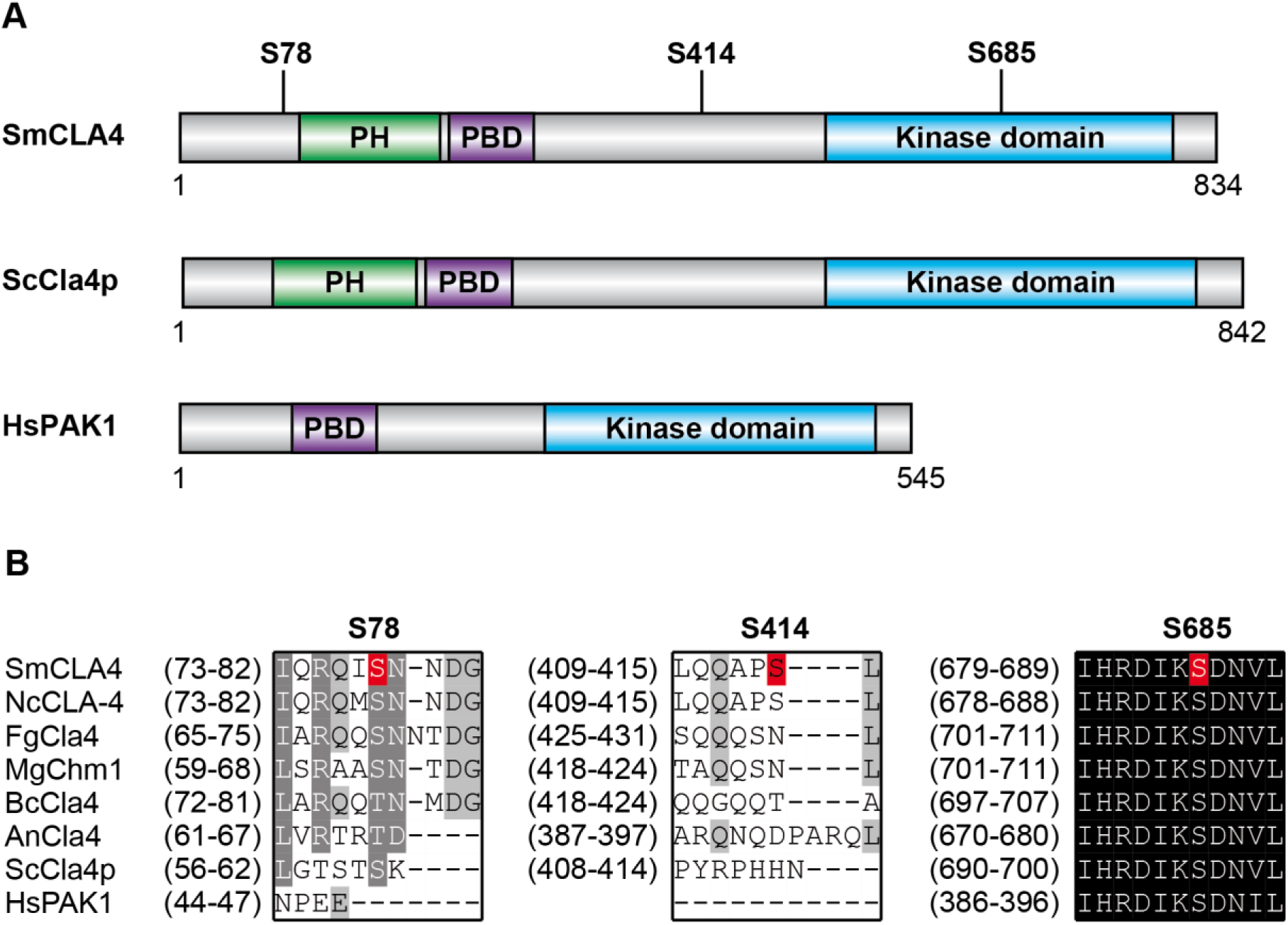
Protein domain structure and identified phosphorylation sites of homologs of CLA4. (A) Primary structure of CLA4 homologs from *S. macrospora* (SmCla4), *S. cerevisiae* (ScCla4p), and humans (HmPAK1). Numbers above the SmCla4 protein indicate residues, representing phosphorylation sites (B) Comparison of phosphorylated regions from *S. macrospora* (Sm) CLA4 with other CLA4-like proteins from *N. crassa* (Nc), *F. graminaerum* (Fg), *Magnaporthe grisea* (Mg), *B. cinerea* (Bc), *A. nidulans* (An), *S. cerevisiae* (Sc), and *H. sapiens* (Hs). Phosphorylation site of CLA4 from *S. macrospora* CLA4 are highlighted in red, which were detected in this investigation. Black color indicates highly conserved amino acid residues. Abbreviations: PBD, p21-binding domain; pleckstrin homology (PH) domain

For functional characterization of CLA4 in *S. macrospora*, a deletion strain was constructed as described in experimental procedures. The homokaryotic Δcla4 strain served for further investigations. As shown in Figure 5A and B, Δcla4 forms ascogonia, protoperithecia, and perithecia within 7 days. However, the perithecia of Δcla4 have a shortened neck and are therefore smaller than wild-type perithecia, even after 14 days of growth (Fig. 5B). Moreover, less than 10% mature asci compared to wt are generated. Δcla4 shows swelling and hyper-branching in vegetative hyphae as well as a very dense growth (Fig. 5C, Fig. S5). Vegetative growth tests showed that Δcla4 displays a strongly reduced growth rate of 3 mm/d in comparison to the wild-type with 28.11 ± 0.79 mm/d on Sordaria Westergaards medium (SWG) (Fig. 6). Previously, we have reported that all STRIPAK mutants have defects in hyphal fusion (Kück et al. 2016). However, in Δcla4, we observed no fusion defect (Fig. S5B). Complementation analysis was performed by transforming plasmid pNA-8843 into the Δcla4 deletion strain. This plasmid carries the *cla4* gene under the control of the native promotor as well as the nourseothricin (*nat*) resistance cassette. After selection of primary transformants on media containing hygromycin B and nourseothricin, ascospore isolates of fertile transformants were generated and phenotypically characterized. The ectopic integration of the *cla4* gene under expressional control of its native promotor completely restored the wild-type phenotype in Δcla4.

**Figure 5:**
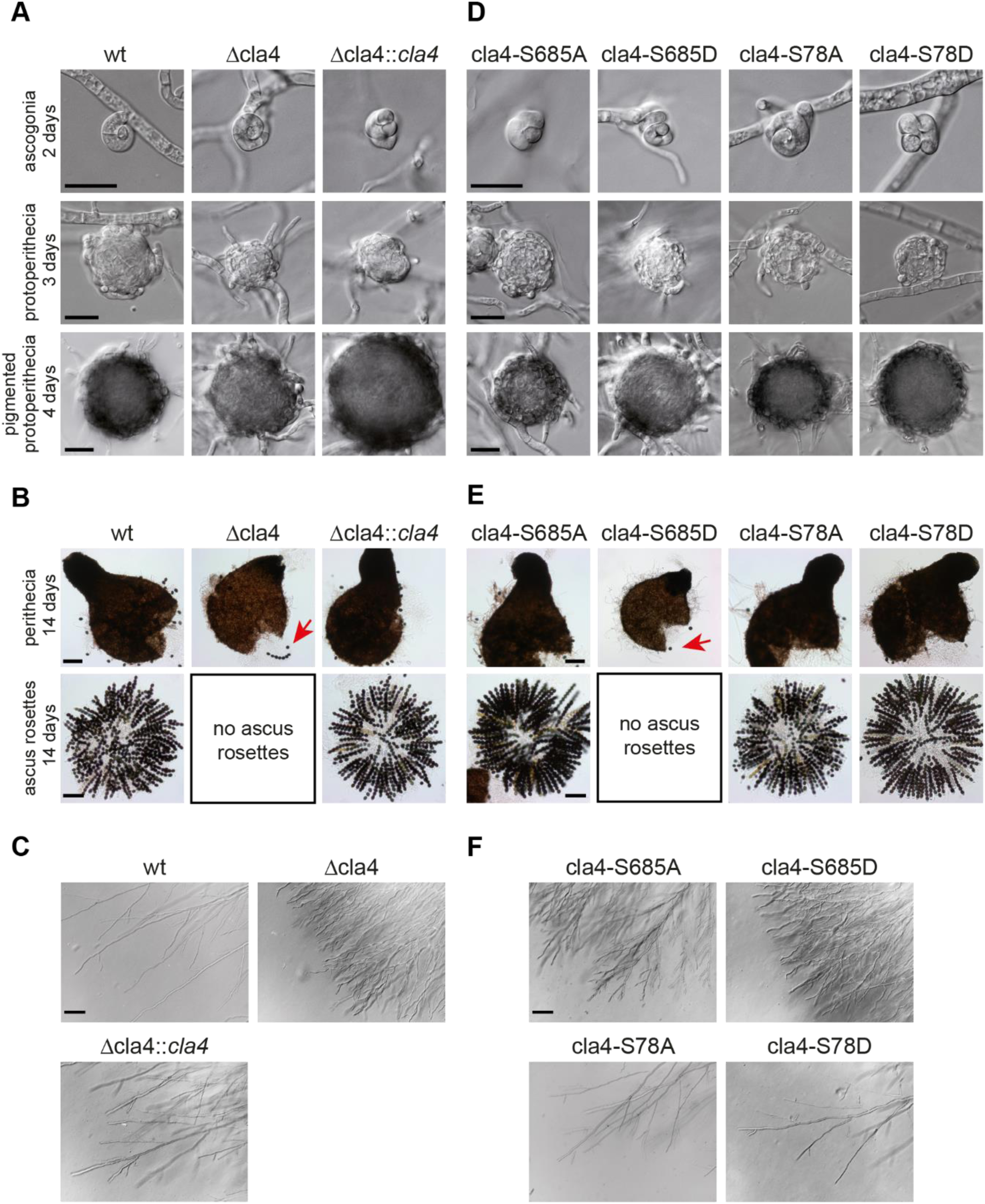
Microscopic analysis of sexual development and hyphal morphology. As indicated, images were obtained from wild type, Δcla4, and transformants, carrying wild type, phospho-mimetic or phosphodeficient versions of *cla4*. All gene constructs were transferred into Δcla4 strain. (A, D) Images of developmental stages of strains grown on BMM-coated slides at 27°C for 2-4 d. Scale bars indicate 20 µm. (B, E) For microscopic analysis of perithecia and ascospores, strains were grown on BMM medium at 27 °C for 14 d. Red arrows indicate ascospores. Scale bar represents 100 µm. (C, F) Documentation of hyphal morphology in the edge region of the colony. Strains were grown on BMM-coated slides at 27°C for 3-7 d. Scale bars indicate 100 µm. In all cases, we show a single representative strain out of at least three individual transformants.

**Figure 6:**
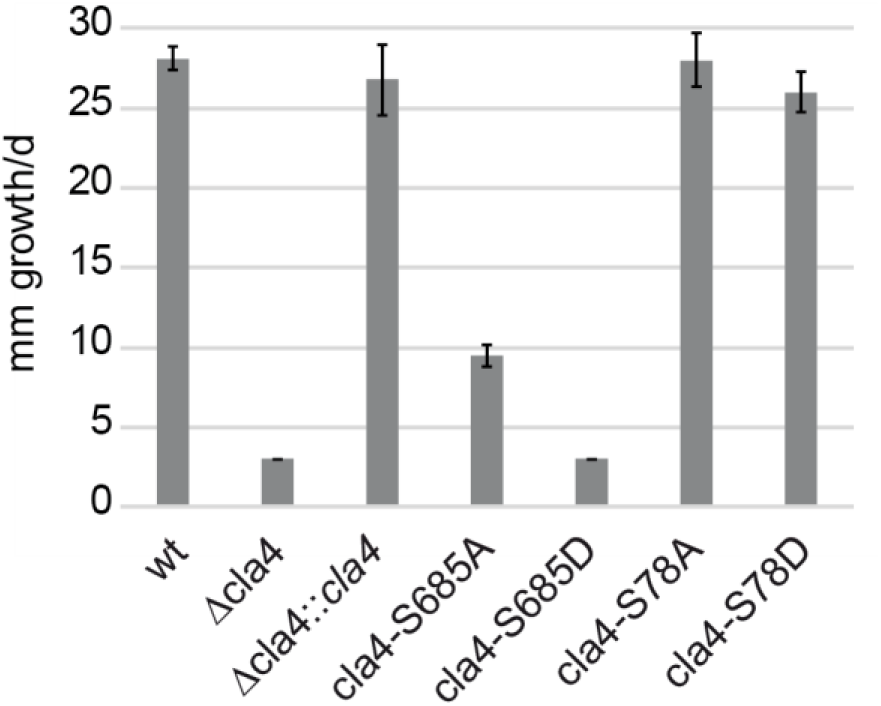
Growth analysis of wild type, Δcla4, and recombinant strains, carrying wild type, phospho-mimetic or phospho-deficient versions of *cla4*. Vegetative growth tests were performed with strains grown in petri dishes on solid SWG medium at 27°C for two days. The graph shows means and standard deviations from three biological replicates per strain. All gene constructs were transferred into Δcla4 strain.

### Phospho-mimetic and -deficient mutants of the PAK kinase CLA4 show defects in sexual development and hyphal growth

In *S. macrospora*, proteomic and phosphoproteomic analyses of the STRIPAK deletion strains Δpro11, Δpro22, and Δpp2Ac1, in comparison to the wild-type identified three phosphorylation sites of the PAK1 homologue CLA4. Two phosphorylation sites, S78 and S414, were found in weakly conserved regions, while S685 is located in the catalytic domain (Fig. 4). As depicted in Table 1, only the phosphorylation site S685 was identified as differentially regulated. The MS data showed more than two-fold upregulation in Δpp2Ac1 and Δpro22 and three-fold upregulation in Δpro11. At the same time, the overall level of CLA4 remained stable in all strains, further strengthening the hypothesis of the STRIPAK complex directly acting on those phosphorylation sites. To unravel the role of phosphorylation site S685 in fungal development, related phospho-mimetic and -deficient variants of CLA4 were generated (Fig. 7). For this purpose, single base pairs of the triplet encoding S685 were mutated, resulting in substitution to either alanine (blocks phosphorylation) or aspartic acid (mimics phosphorylation because of a negative charge) in the derived protein. Mutation of the phosphorylation site S78 to A78 or D78 served as control since it is located in a weakly conserved region.

**Figure 7:**
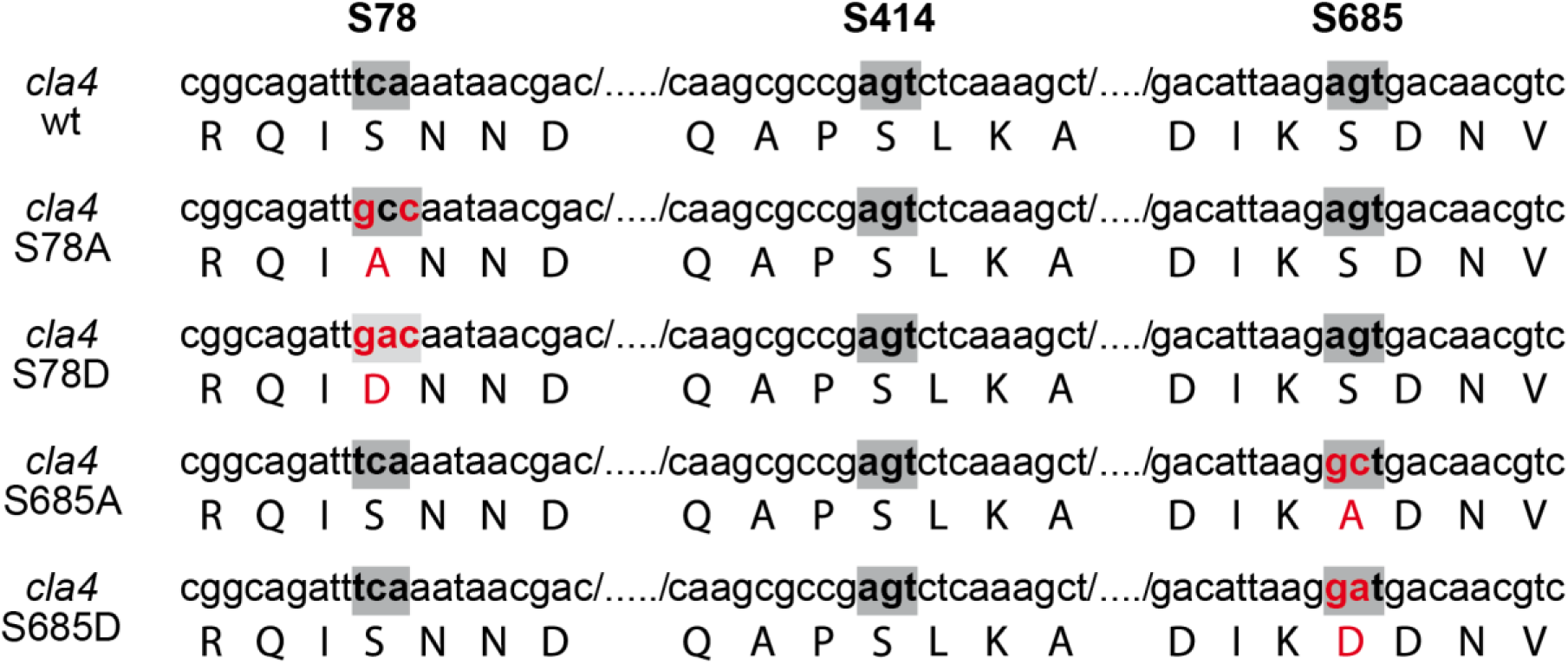
Phospho-mimetic and –deficient versions of *cla4* used for mutagenesis studies. Lower case and capital letters indicate coding sequence of *cla4* and the derived amino acid sequence in proximity to serine phosphorylation sites S78, S414 or S685. The base pair triplets encoding the phosphorylated amino acids are displayed in grey. Red letters mark single base pair substitutions and the corresponding amino acid substitutions S78A, S78D, S685A and S685D.

The four plasmids pNA-8843-S78A, pNA-8843-S78D, pNA-8843-S685A, and pNA-8843-S685D, encoding the phospho-mimetic and -deficient variants of CLA4 were transformed into the Δcla4 deletion strain. After selection of primary transformants by hygromycin B and nourseothricin resistance, ascospore isolates of fertile transformants (cla4-S78A, cla4-S78D, cla4-S685A, and cla4-S685D) were generated and phenotypically analyzed. Representative isolates of all mutants are shown in Figs. 5 and 6.

Sexual development and hyphal growth were investigated in all selected strains and compared to wild type, Δcla4, as well as Δcla4::*cla4*. As depicted in Figure 5D,E, the analysis of different developmental stages revealed that Δcla4::*cla4*, cla4-S78A, and cla4-S78D form wild-type-like ascogonia, protoperithecia, as well as pear-shaped perithecia. Further, vegetative growth of all these strain is identical to the wild-type (Fig. 6). In contrast, cla4-S685A shows swelling of hyphae, hyperbranching, and a reduced growth rate, albeit sexual development seems to be wild type-like (Fig. 5 and 6). Strikingly, a completely different phenotype was observed for cla4-S685D. In detail, this strain generates deformed perithecia and the number of discharged ascospores is highly reduced. Moreover, the phenotype is similar to that of Δcla4. Of note is that this phenotype is also evident when we investigated the morphology of vegetative hyphae (Fig. 5F, Fig. S5).

## Discussion

The conserved STRIPAK complexes from fungi, animal, or humans have the capacity to phosphorylate or dephosphorylate target proteins. The complex interacts with other conserved signalling complexes, which might be potential dephosphorylation targets (Hwang and Pallas 2014; Kück et al. 2016). Although some substrates of STRIPAK have been identified, the picture is still far from complete. Thus, to identify targets of STRIPAK likely involved in fungal development, we compared the proteome and phosphoproteome of wild-type and three STRIPAK mutants. Here, we used quantitative phosphoproteomics to identify potential targets of STRIPAK.

### Quantitative phosphoproteomic analysis in fungi

iTRAQ-based phosphoproteomic analysis has already been performed in diverse filamentous fungi such as *Aspergillus fumigatus* (Cagas et al. 2011; Adav et al. 2015), *Beauveria bassiana* (Wang et al. 2016), *Fusarium graminearum* (Taylor et al. 2008), and *Neurospora crassa* (Jonkers et al. 2014; Xiong et al. 2014). In this work, we present an in-depth quantitative phosphoproteomic data set of *S. macrospora* in conjunction with corresponding global proteome analysis, allowing for differentiation between changes in total protein expression and true alterations of specific phosphorylation levels. With the identification and quantification of 8,908 phosphorylation sites across all conditions, our data set exhibits substantially higher coverage than comparable work with other filamentous fungi to date (Jonkers et al. 2014; Xiong et al. 2014; Franck et al. 2015; Zhou et al. 2019). Here, we provide the first comprehensive STRIPAK-dependent quantitative proteome and phosphoproteome analyses in a eukaryotic organism. Further, from the sum of all quantitative data, we substantially extend the number of potential STRIPAK targets. Finally, this study also contributes significantly to our current understanding of the mechanistic function of STRIPAK as a phosphatase and kinase signalling complex.

### Phosphorylation of the potential STRIPAK target CLA4 controls fungal development

One of the putative potential STRIPAK targets is the PAK kinase CLA4, which was identified with three different phosphorylation sites. The phenotypical analysis of a Δcla4 deletion strain revealed that CLA4 is involved in sexual development. This finding is similar to previous reports that have investigated the function of CLA4 homologues in other filamentous fungi. For example, in *Bipolaris maydis*, the CLA4 homologue regulates the formation of fruiting bodies as well as ascospore development (Kitade et al. 2019). Similarly, CLA4 homologues in *F. graminaerum*, *N. crassa*, and *M. grisea* affect ascospore development and release, but not fruiting body morphology (Li et al. 2004; Park et al. 2011; Wang et al. 2011).

Further, Δcla4 exhibits dense growth and altered branching in vegetative hyphae, suggesting that CLA4 is involved in regulating cell polarity in *S. macrospora*. This observation is consistent to reports by others, thus indicating that CLA4 function is conserved in filamentous ascomycetes (Rolke and Tudzynski 2008; Lichius et al. 2014; Tian et al. 2015; Kitade et al. 2019). The branching pattern of hyphae might be due to the non-regular distribution of the Spitzenkörper, as was suggested for a *B. maydis* Δcla4 deletion strain (Kitade et al. 2019). In *S. cerevisiae*, the PAK1 homologue Cla4p has also been extensively investigated. Several studies indicate that Cla4p regulates important biological processes, including cell polarity, cell cycle progression, cytokinesis, and gene transcription (Cvrcková et al. 1995; Benton et al. 1997; Lin et al. 2009). In addition, overexpression of Cla4p was shown to affect the pheromone-induced cell cycle arrest. Thus, Cla4p has the potential to control the PR pathway (Heinrich et al. 2007). Taken together, these studies support the view that CLA4 homologues play a significant role in fungal development.

The human CLA4 homologue PAK1 is the best studied member of the PAK family and is involved in a variety of cellular processes such as cytoskeleton remodelling, cell cycle control, the DNA damage response, cell motility, cell apoptosis, and neurodevelopment. Several studies in human showed that PAK1 is crucial for cardiac excitation, and muscle contraction dynamics by regulation of ion channel activity (Kumar et al. 2017; Wang et al. 2018). Further, PAK1 is upregulated and activated in several human tumor types, including breast, colon, and brain tumors (Kumar et al. 2006). PAK1, a member of group A PAK kinases, is regulated by an auto-inhibition mechanism. In an inactivated state, the autoinhibitory domain (AID) overlaps with the PBD domain and binds the catalytic domain of another PAK1 protein, resulting in the formation of a homodimer. The binding of the small GTPases CDC42 or RAC to the PBD domain leads to a conformational change in the AID, followed by dissociation from the catalytic domain of the other PAK1 molecule. Both PAK proteins become auto-phosphorylated, and therefore, activated at several sites, including the activation loop within the kinase domain (Zhao and Manser 2012; Kumar et al. 2017). In human as well as yeast, various phosphorylation sites of PAK1 and Cla4p have been detected and experimentally verified. Further, the generation of phospho-mimetic and -deficient variants of PAK1 regarding numerous phosphorylation sites, including S144, S223, and T423, revealed their importance for kinase activity (Chong et al. 2001; Ng et al. 2010; Shin et al. 2013). Since PAK1 is associated with several disease phenotypes (Kumar et al. 2017; Wang et al. 2018), PAK1 emerged as a therapeutic target and several PAK1 inhibitors were developed (Kumar et al. 2017).

Our phosphoproteomic analysis of the STRIPAK deletion strains and the wild-type revealed three different phosphorylation sites. S78 and S414 are located in weakly conserved regions of the protein, while S685 locates in the highly conserved catalytic domain. Only phosphorylation site S685, which has neither been characterized in human nor yeast, was regulated in the STRIPAK deletion strains. The phospho-deficient mutation of this site leads to reduced growth as well as slightly enhanced swelling and hyper-branching in vegetative hyphae, but does not affect the formation of fruiting bodies and ascospore development. These results indicate that phosphorylation is important for the functional mechanism of CLA4 in vegetative hyphal growth. In contrast, the phospho-mimetic mutant S685D resulted in severe defects of sexual development and vegetative hyphal growth comparable to the Δcla4 deletion strain, thereby suggesting that dephosphorylation of S685 might be even more critical for proper CLA4 function. Thus, phosphorylation of S685 from CLA4, a member of the conserved p21-activated kinase (PAK) family, seems to be STRIPAK dependent. Previous studies in human already described this amino acid as a potential substrate-binding site of PAK1 (Ng et al. 2010), thus emphasizing the significance of the phosphorylation site S685.

### CLA4 plays an important role in connecting the STRIPAK complex

From our analysis, we conclude that CLA4 is a putative dephosphorylation target of the STRIPAK complex, and AP-MS analysis indicates further an association with the striatin-interacting protein PRO22 (Märker 2019). Previous studies in human showed that the association of the CLA4 homologue PAK1 with PP2A plays a role in regulating Ca^2+^ homeostasis in the heart. In particular, these studies indicated that PAK1 induces auto-dephosphorylation of the catalytic subunit PP2Ac at the phosphorylation site Y307 through a scaffolding mechanism. Subsequently, PP2A antagonizes the effects of cAMP-dependent kinase (PKA) on Ca^2+^ channels activity (Ke et al. 2008; Ke et al. 2013). Further, there is evidence that PAK1 and PP2A form a signalling module in mast cells, where PP2A dephosphorylates PAK1 at T423 in the activation loop, resulting in the disassembly of PP2A (Staser et al. 2013).

In conclusion, our study indicates that the STRIPAK complex regulates numerous signalling pathways controlling different biological processes in *Sordaria macrospora*. These results increase our current basic knowledge about the function of the eukaryotic STRIPAK complex and further support the notion that STRIPAK is a central regulator of diverse cellular processes. Moreover, our study provides clear evidence that phosphorylation and dephosphorylation of residue S685 of Cla4 is crucial for fungal cellular development. Finally, since this amino acid has not yet been reported to be phosphorylated in other eukaryotes, our findings will encourage future investigations of this phosphorylated site in CLA4 homologues, which in humans were considered to be therapeutic targets in cancer and allergen-induced disorders (Kichina et al. 2010; Pandolfi et al. 2015).

## Experimental procedures

### Strains and growth conditions

All *S. macrospora* strains (Table S1) were grown under standard conditions, unless otherwise described (Kamerewerd et al. 2008). DNA-mediated transformation was performed as described previously (Nordzieke et al. 2015), but lacking caylase treatment during protoplast formation. Transformants were selected on medium containing hygromycin B (80 U/ml) and/or nourseothricin (100 µg/ml). Growth tests were performed three times with three technical replicates per strain, and hyphal growth was measured as described before (Teichert et al. 2017). For each experiment, petri dishes with solid SWG medium were inoculated with an 8-mm-diameter agar plug and incubated for 2 days. Isolation of DNA and Southern hybridization were performed as described previously (Kamerewerd et al. 2008).

For cloning and propagation of recombinant plasmids, *E. coli* XL1-Blue MRF‘ (Jerpseth et al. 1992) and NEB5α (New England Biolabs, Frankfurt, Germany) were used under standard laboratory conditions (Sambrook and Russell 2001). For plasmid construction, homologous recombination in *S. cerevisiae* PJ69-4a was performed (James et al. 1996) as described previously (Colot et al. 2006). Recombinant yeast strains were selected on minimal medium, lacking uracil; all other experiments with yeast were carried out according to standard protocols (Clontech Yeast Protocol Handbook, PT3024-1).

### Protein extraction, enrichment, and fractionation

For protein extraction, *S. macrospora* strains were precultured in petri dishes with 20 ml liquid BMM with two biological replicates per strain at 27°C for 2 days. Three standardized inoculates of each BMM preculture were transferred in petri dishes with 20 ml liquid BMM and grown at 27°C and 40 rpm for 3 days. For cell wall lysis and protein extraction, mycelium was harvested and freezed within 60 s. The frozen mycelium was ground in liquid nitrogen and suspended in FLAG extraction buffer (50 mM Tris-HCl pH 7.4, 250 mM NaCl, 10% glycerol, 0.05% NP-40,1 mM PMSF, 0.2% protease inhibitor cocktail IV, 1.3 mM benzamidine, as well as 1% phosphatase inhibitor cocktails II and III). Afterwards, the samples were centrifuged at 4°C and 15,000 rpm for 30 min (modified after Teichert et al. 2014). The same lysates were used for proteomic and phosphoproteomic analyses. Protein concentration in all lysates was determined by a calorimetric bicinchoninic acid assay (Pierce BCA protein concentration assay kit) following the manufacturer’s protocol. Carbamidomethylation was performed by reduction of free cysteine residues by the addition of dithiothreitol (DTT) to a final concentration of 10 mM and incubation for 30 min at 56°C followed by alkylation with 30 mM iodoacetamide (IAA) for 30 min at room temperature in the dark. Excess of IAA was quenched by further addition of 10 mM fresh DTT. Samples were purified prior to digestion by ethanol precipitation and resuspended with 40 µl of 6 M guanidinium hydrochloride, followed by dilution with ammonium bicarbonate buffer (pH 7.8) to a final concentration of 0.2 M and addition of CaCl_2_ to a final concentration of 2 mM. Trypsin was added at a 1:20 (protease:substrate (w/w)) ratio and samples were incubated for 14 h at 37°C. Digestion was stopped by addition of 10% Trifluoroacetic acid (TFA) to a final concentration of 1%. Acidified peptides were desalted and quality controlled as described previously (Burkhart et al. 2011). Peptides were then dried down completely using a SpeedVac and resuspended in 0.5M triethylammonium bicarbonate (pH 8.5). Per sample, 150 μg of tryptic peptide were labelled with iTRAQ 8-plex reagents (AB Sciex, Darmstadt, Germany) following the manufacturers protocol. After pooling and quenching, an aliquot corresponding to 70 μg of total peptide amount was taken for global proteome analysis. From this, 35 μg were fractionated by high-pH reversed phase chromatography using an Ultimate 3000 HPLC (Thermo Scientific) equipped with a C18 column (BioBasic-18, 5 μm particle size, 300 Å pore size, 150 x 0.5 mm). A total of 20 fractions was collected in 1 min windows using a concatenated collection mode.

For protein enrichment, the remaining multiplexed sample (1,130 μg) was dried under vacuum and subjected to a phosphopeptide-enrichment protocol using titanium dioxide (TiO_2_, Titansphere TiO, 5 µm particle size, GL Sciences Inc, Japan) adapted from a previous report (Engholm-Keller et al. 2012). For protein fractionation, phosphopeptides were fractionated by hydrophilic interaction liquid chromatography on an Ultimate 3000 HPLC (Thermo Scientific, Dreieich, Germany) and a total of 13 fractions was collected.

### LC-MS/MS analysis

Samples for global proteome analysis were subjected to LC-MS/MS analysis using an Ultimate 3000 nanoRSLC HPLC coupled to a Q Exactive HF mass spectrometer (both Thermo Scientific). Preconcentration of peptides was performed on a precolumn (Pepmap RSLC, Thermo Scientific, C18, 100 µm x 2 cm) for 10 min at 20 µl/min flow (0.1% TFA) followed by separation on a 75 µm x 2 cm C18 main column (Pepmap RSLC, Thermo Scientific). A binary gradient was used with 0.1% formic acid (FA) as solvent A and 84% acetonitrile (ACN), 0.1 % FA as solvent B. A linear gradient was used with solvent B increasing from 3 to 35% in 120 min, The MS was operated in the data-dependent acquisition (DDA) mode, first performing a survey scan from 300 to 1,500 m/z at a resolution of 60,000, with the AGC target set to 3 x 10^6^ and the maximum injection time to 120 ms. The polysiloxane ion at m/z 371.101236 was used as lock mass and the top 15 most intense ions were subjected to higher energy collisional dissociation (HCD) and subsequent MS/MS analysis. HCD normalized collision energy was set to 31% and quadrupole isolation width was limited to 0.7 m/z. MS/MS scans were acquired at a resolution of 15,000 with an AGC target value of 2 x 10^5^, a maximum allowed injection time of 250 ms, and a fixed first mass of 90 m/z. Dynamic exclusion of precursor ions was set to 30 s. To reduce charge states of precursor ions, a 10% (v/v) NH_4_OH solution was placed at the nano source as described previously (Thingholm et al. 2010). Phosphopeptide fractions were subjected to LC-MS/MS analysis using an Ultimate 3000 nanoRSLC HPLC coupled to an Orbitrap Fusion Lumos MS (both Thermo Scientific). LC separation was performed as described above, using a gradient from 3 to 42% B in 90 min. The MS was operated in the DDA mode with a 3 s cycle time (Top Speed). Survey scans were acquired at a resolution of 120,000 while for MS/MS scans this was set to 30,000. AGC target values were 2 x 10^5^ and 1 x 10^5^, respectively. Maximum injection times were 50 ms and 200 ms and HCD collision energy was 40%. Quadrupole isolation was performed with a 0.8 m/z window and all other settings were kept as described above.

### Proteomics data analysis

Acquired raw data was analysed with Proteome Discoverer 1.4 (Thermo Scientific), incorporating the search algorithms Mascot (Version 2.4.1, Matrix Science), Sequest HT, and MS Amanda. The searches were conducted in a target/decoy manner against a *S. macrospora* protein sequence database (10,091 target sequences) with the same settings for all three algorithms: Precursor mass tolerance was set to 10 ppm and fragment mass tolerance to 0.02 Da. Cleavage specificity was set to trypsin with a maximum of 2 allowed missed cleavages. iTRAQ-8 plex on peptide N-termini and lysines as well as carbamidomethylation of cysteines was set as fixed modification, while oxidation of methionine and phosphorylation (only phosphoproteome data) of serine, threonine, or tyrosine was allowed as variable modifications. For phosphoproteome analysis, the phosphoRS (version 3.1) node (Taus et al. 2011) was used to determine modification site confidence. Percolator was used to filter the results to a false discovery rate (FDR) of 1% on the PSM level and only rank 1 hits were allowed. For global proteome data, a minimum of 2 uniquely identified peptides per protein were required, while for the phosphoproteome data, only quantified PSMs with identified phosphorylations and a phosphoRS site probability ≥ 90 % were exported.

To correct for systematic errors during sample labelling, global proteome data was normalized by correction factors calculated from the summed total intensities of all iTRAQ channels. Mean ratios of biological replicates were calculated and proteins with a change in abundance greater than two times the standard deviation of the respective condition were considered as regulated. For analysis of phosphoproteome data, an Excel macro was used provided by Taus et al. (2011). Data were normalized using the correction factors determined from the global data. Only ratios of confidently localized phosphorylations were used in the analysis and mean of biological replicates was calculated. Two times the standard deviation of the total data set of the respective condition was used as the criterion to determine regulation of each phosphopeptide.

### Phosphorylation motif analysis

To identify overrepresented consensus motifs of the identified phosphorylation sites, seven flanking amino acids up- and downstream of the modified residues were extracted. The motifs of up- or downregulated sites in the individual deletion strains were uploaded to the MoMo web server (Cheng et al. 2018). Significantly enriched motifs were identified using the motif-x algorithm and the *S. macrospora* protein database (10,091 sequences) as context sequence and requiring a minimum number of 20 occurrences and a p-value of threshold of 1E-6.

### Construction of plasmids

All oligonucleotides and plasmids are listed in the supporting information (Tables S2 and S3). The knock-out plasmid pKO-8843 for the deletion of *cla4* was generated by yeast recombination. For this purpose, the 5’- and 3’-flanking regions of *cla4* were amplified by PCR from genomic DNA of *S. macrospora* with the primers 8843-5fw/8843-5rv-hph and 8843-3fw/8843-3rv, respectively, and transformed together with an *hph* cassette cut with *Eco*RI out of plasmid pDrive-hph (Nowrousian and Cebula 2005) and *Eco*RI/*Xho*I-linearized plasmid pRS426 (Christianson et al. 1992) in yeast.

Yeast recombination was also used to obtain the complementation plasmid pNA-8843. For this purpose, the *cla4* gene and the corresponding 5‘- and 3‘-flanking regions containing the promotor and terminator of *cla4* were amplified from genomic DNA of *S. macrospora* with the primers 8843-5fw and 8843-3rv, and recombined into the *Hind*III/*Xho*I-linearized plasmid pRSnat (Klix et al. 2010).

For construction of plasmids encoding phospho-mimetic and -deficient variants of CLA4 with the amino acid substitutions S685A or S685D, the Q5® Site-Directed Mutagenesis Kit (New England Biolabs) was used according to the manufacturer’s protocol. For the PCR, pNA-8843 was used as template with the primers Cla4-S685A-2-fw/Cla4-S685-2-rv and Cla4-S685D-2-fw/Cla4-S685-2-rv, resulting in pNA-8843-S685A and pNA-8843-S685D, respectively.

Plasmids encoding the phospho-mimetic and -deficient variants of CLA4 with the amino acid substitutions S78A or S78D were generated by the yeast recombination system. PCRs with the primers 8843-5fw-NheI/8843-S78A-rv, 8843-5fw-NheI/8843-S78D-rv, 8843-S78A-fw/8843-rv, and 8843-S78D-fw/8843-rv were performed using genomic DNA of *S. macrospora* as template. Using primers 8843-3fw/8843-3rv-NheI, the 3’ end of cla4 was amplified. All PCR fragments, the *Eco*RI-linearized *hph* cassette, (pDrive-hph), together with the *Eco*RI/*Xho*I-linearized yeast vector pRS426 were transformed into yeast (PJ69-4a). The resulting plasmids carry the mutated *cla4* gene and received the designations pCla4-S78A and pCla4-S78D. Plasmids pCla4-S78A and pCla4-S78D were restricted with *Stu*I/*Ksp*I, and fragments carrying the mutated *cla4* gene were integrated into *StuI*/*Ksp*I-linearized pNA-8843. The resulting plasmids were designated pNA-8843-S78A and pNA-8843-S78D, respectively.

### Generation of Δcla4, and phospho-mimetic and -deficient mutants

For generation of a Δcla4 strain, plasmid pKO-8843 was linearized with *Eco*RI and transformed into Δku70 (Pöggeler and Kück 2006). The primary transformants were selected for hygromycin B resistance and verified by PCR analysis (data not shown). Ascospore isolates of the Δcla4 strain with the wild-type genetic background (lacking the Δku70 mutation) were obtained by mating with spore colour mutant fus as described previously (Kück et al. 2009; Nowrousian et al. 2012). The recombinant strains were verified by PCR and Southern blot analysis (Fig. S4). To restore the wild-type phenotype, plasmid pNA-8843 was transformed into the Δcla4 recipient. The corresponding transformant was designated Δcla4::*cla4*.

Transformation of pNA-8843-S78A, pNA-8843-S78D, pNA-8843-S685A, and pNA-8843-S685D into the Δcla4 recipient strain resulted in cla4-S78A, cla4-S78D, cla4-S685A, and cla4-S685D carrying the phospho-mimetic and -deficient mutations in the *cla4* gene. The point mutations of *cla4* in diverse transformants were verified by PCR and DNA sequencing analysis.

### Microscopic investigations

Light microscopy of different developmental stages and hyphal fusion events was performed by differential interference contrast (DIC) microscopy using an AxioImager microscope (Zeiss) with a Photometrix Cool SnapHQ camera (Roper Scientific) and the software MetaMorph (version 7.7, Universal Imaging). For light microscopy of smashed perithecia and ascospores, the AxioPhot microscope (Zeiss) was used, and images were obtained with an AxioCam color and ZEN software (version 2.3, Zeiss). For microscopic investigation of different developmental stages, *S. macrospora* strains were grown on BMM-covered slides at 27 °C and constant light for 2-7 d (Engh et al. 2007). For microscopic analysis of hyphal fusion events, strains were incubated on cellophane-covered solid MMS medium at 27°C for 2-4 d (Rech et al. 2007). For microscopic analysis of perithecia and ascospores, strains were grown at 27 °C and constant light for 14 d. After transferring perithecia to slides, 0,96 % NaCl solution was added, and perithecia were smashed open.

### Data availability

The mass spectrometry proteomics data have been deposited to the ProteomeXchange Consortium via the PRIDE partner repository (Vizcaíno et al. 2014) with the dataset identifier PXD014858.

## Acknowledgements

We thank Ingeborg Godehardt for excellent technical support, and Dr. Ines Teichert for helpful and critical discussion. This study was funded by the Deutsche Forschungsgemeinschaft (Bonn, Germany) (KU517/16-1, KU517/16-2, SI835/6-1, SI835/8-2).

RM, BBL, ABR, AS, UK conceived and designed the study. RM and BBL acquired experimental data. RM, BL, AS and UK analysed and interpreted the data. RM, BBL, ABR and UK wrote the manuscript.

## Supporting information

**Table S1:** S. macrospora strains used in this study.

| Strain | Characteristics | Source/Reference |
| --- | --- | --- |
| R19027 | wild type, fertile | culture collection<sup>1</sup> |
| S70823 | *fus*, spore color mutant, fertile | culture collection<sup>1</sup> |
| S151468 | $\Delta ku70::nat^r$, fertile | culture collection<sup>1</sup> |
| A1844 | $\Delta pp2ac1::hph^r$, sterile | Beier et al. (2016) |
| S5.7 | $\Delta pro11::hph^r$, sterile | Bloemendal et al. (2012) |
| S56 | $\Delta pro22::hph^r$, sterile | Bloemendal et al. (2012) |
| $\Delta cla4$ (**RM1861**,<br>RM1911, RM1918) | $\Delta cla4::hph^r$, fertile | This work |
| $\Delta cla4::cla4$ (**RM2094**,<br>RM2078, RM2088) | $\Delta cla4::hph^r$<br>*cla4(p)::cla4::cla4(t)::nat^r*, fertile | This work |
| *cla4*-S685A<br>(**RM2987**, RM2984,<br>RM3006) | $\Delta cla4::hph^r$<br>*cla4(p)::cla4-S685A::cla4(t)::nat^r*, fertile | This work |
| *cla4*-S685D | $\Delta cla4::hph^r$<br>*cla4(p)::cla4-S685D::cla4(t)::nat^r*, fertile | This work |
| *cla4*-S78A (**RM3315**,<br>RM3327, RM3334) | $\Delta cla4::hph^r$<br>*cla4(p)::cla4-S78A::cla4(t)::nat^r*, fertile | This work |
| *cla4*-S78D (**RM3464**,<br>RM3471, RM3499) | $\Delta cla4::hph^r$<br>*cla4(p)::cla4-S78D::cla4(t)::nat^r*, fertile | This work |

Numbers in brackets indicate individual fungal ascospore isolates. Bold printed strains mark representative isolates shown in the results section; *nat<sup>r</sup>*, nourseothricin resistance gene; *hph<sup>r</sup>*, hygromycin B resistance gene; <sup>1</sup> Department of General and Molecular Botany, Ruhr-University Bochum, Germany

**Table S2:**
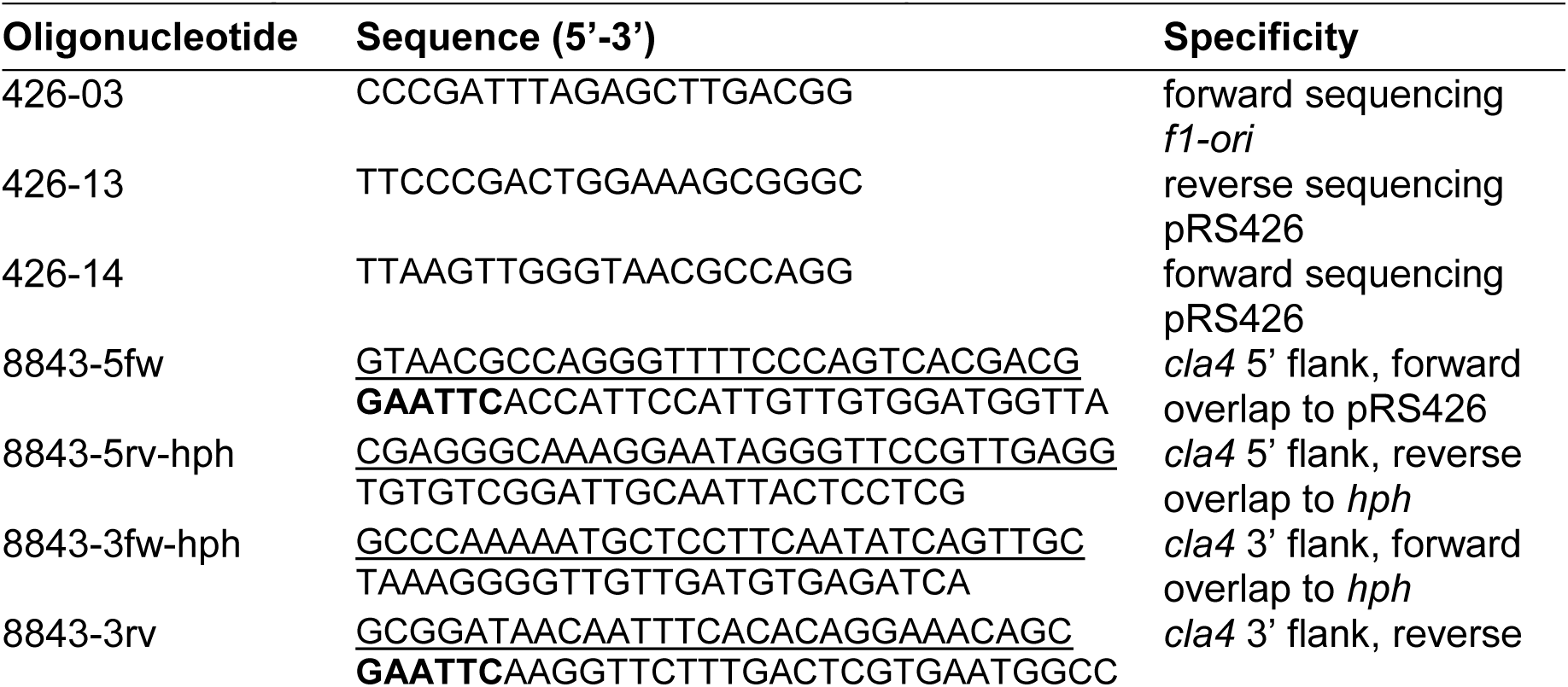
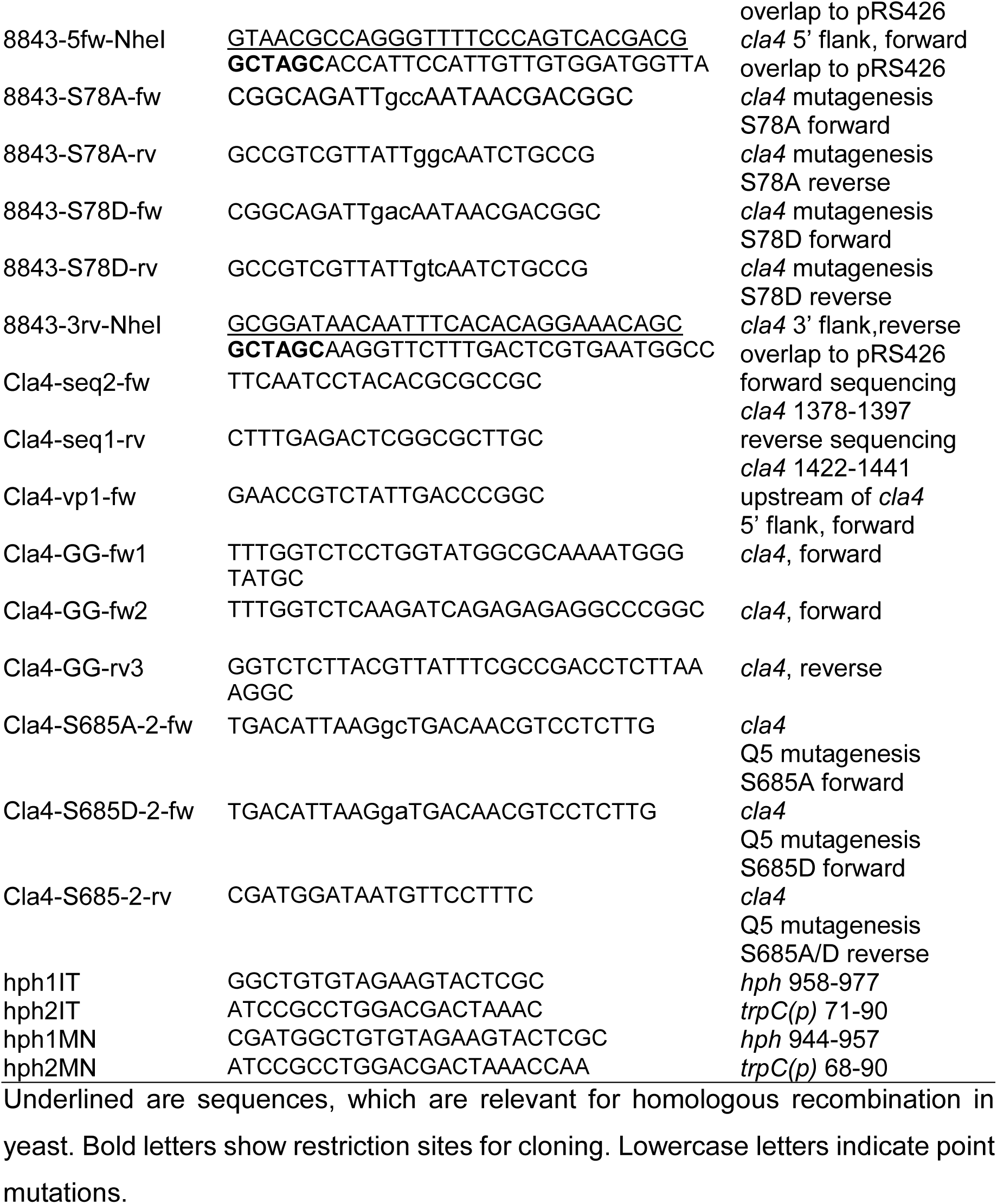
Oligonucleotides used in this study.

**Table S3:**
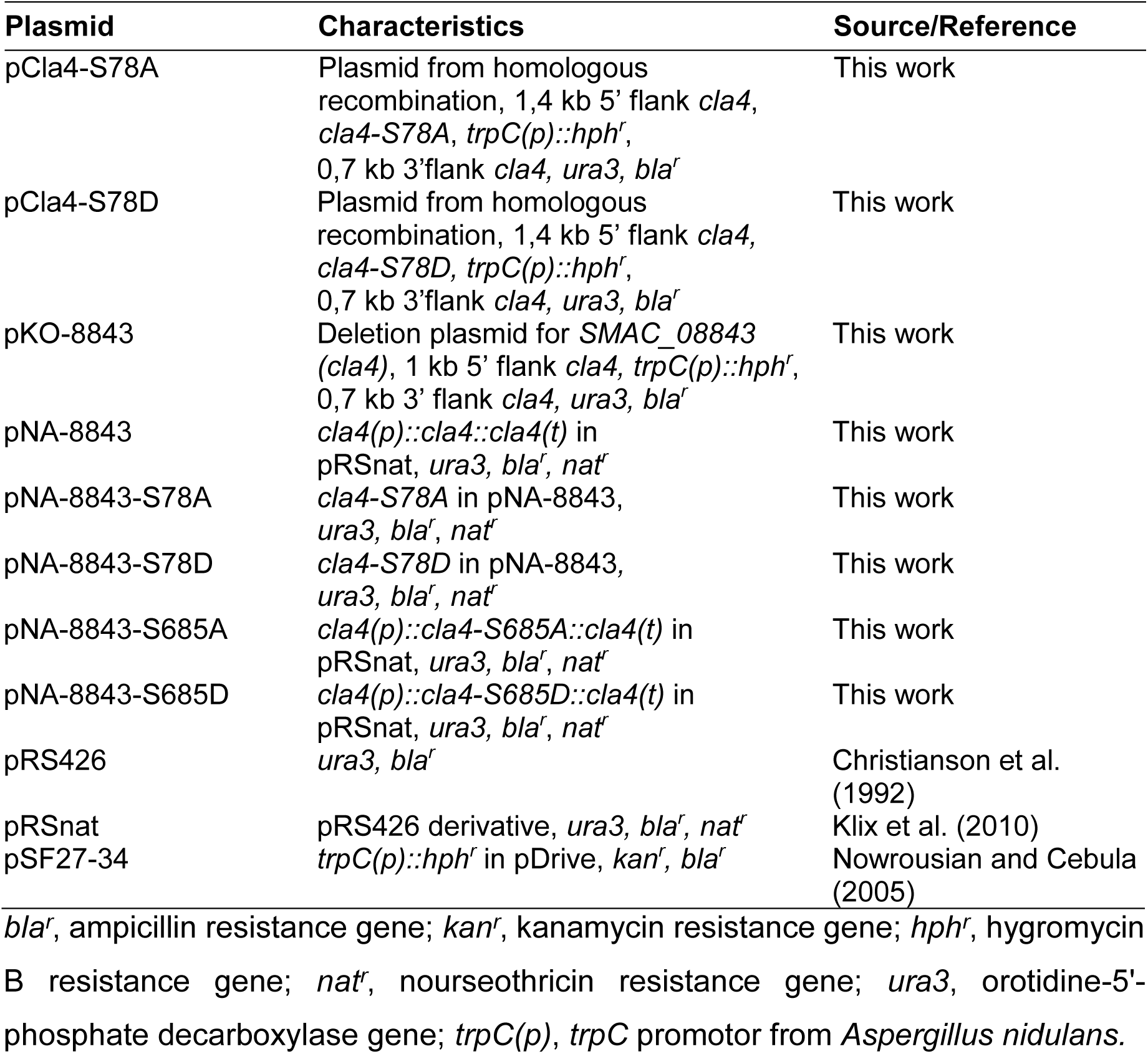
Plasmids used in this study.

**Figure S1:**
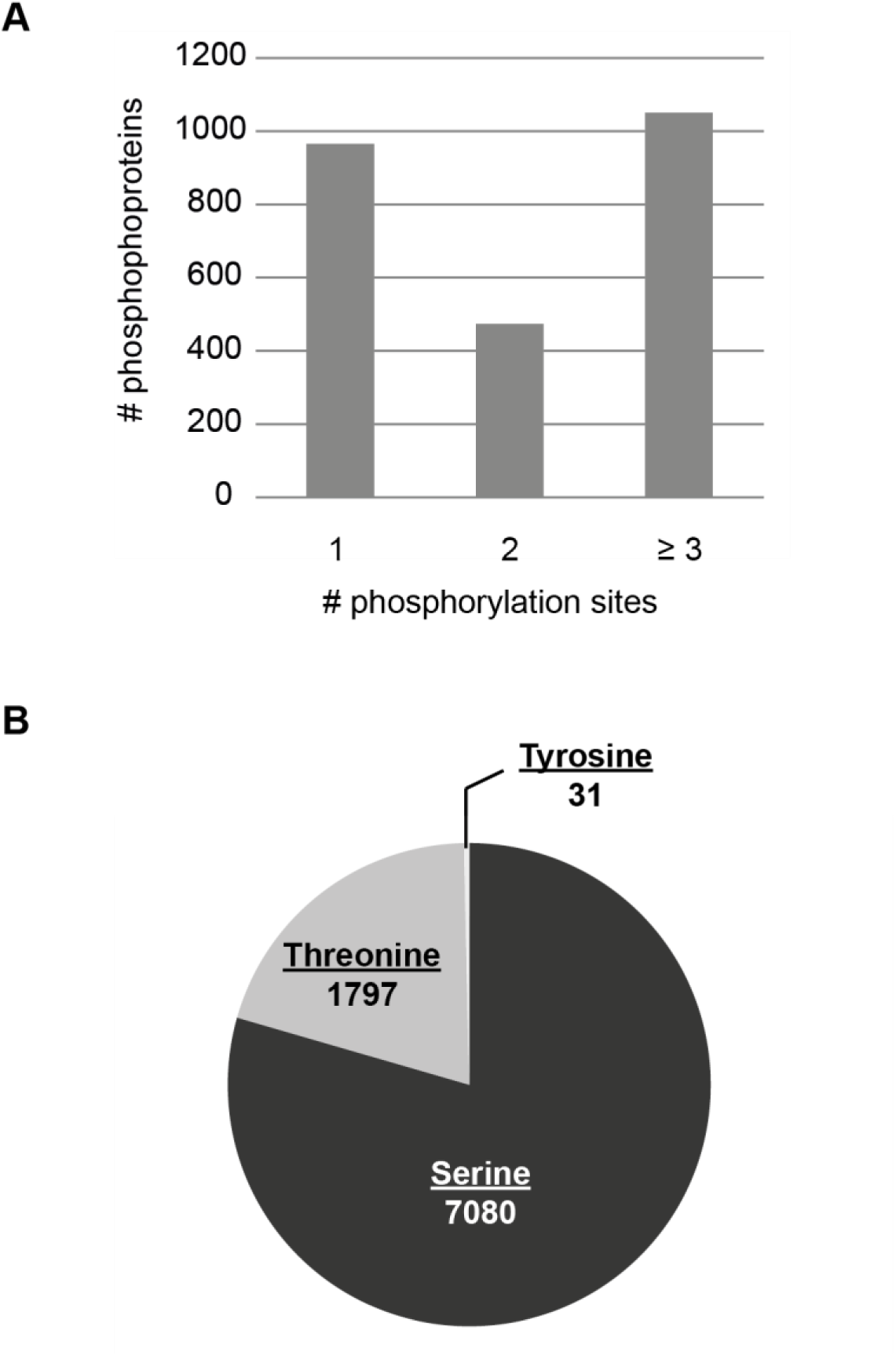
Analysis of identified phosphorylation sites in wild type, Δpp2Ac1, Δpro11 and Δpro22. (A) Absolute numbers of phosphoproteins with one, two, three or more phosphorylation sites are shown. (B) Pie chart displays the identified numbers of phosphorylated serine, threonine, and tyrosine residues.

**Figure S2:**
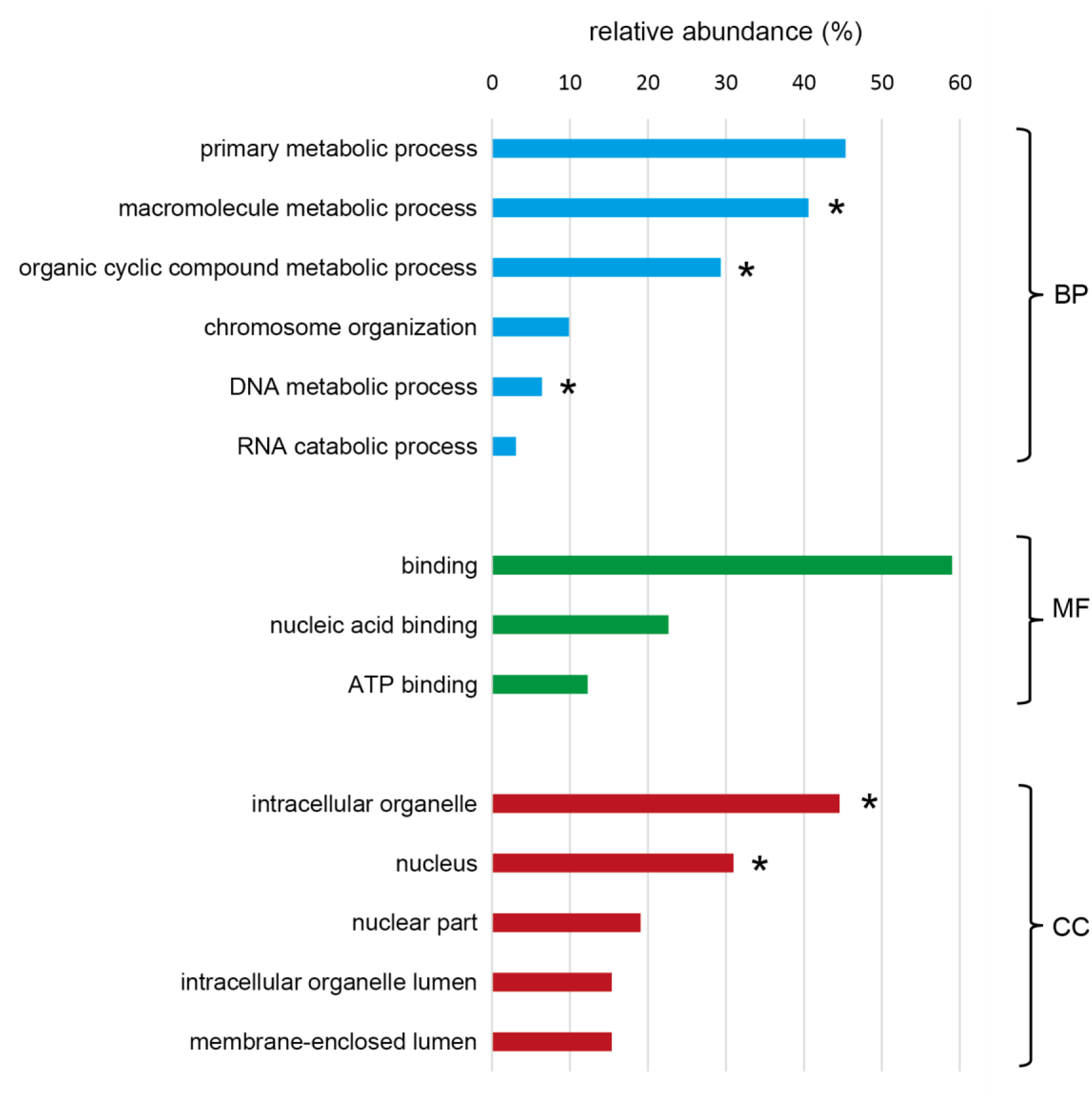
GO enrichment analysis of proteins with regulated phosphorylation sites in STRIPAK deletion strains. Phosphoproteomic analysis identified 781 phosphoproteins, which exhibit regulated phosphorylation sites in Δpp2Ac1, Δpro11, and Δpro22 relative to the wild type. GO enrichment analysis of these proteins performed with the ontologizer 2.0 command line tool revealed the overrepresented GO terms in the categories of biological process (BP), molecular function (MF), and cellular component (CC). GO terms are sorted by percentage of proteins among all 781 phosphoproteins. All GO terms exhibit a p-value ≤ 0,05. Asterisks indicate GO terms with a p-value ≤ 0,01.

**Figure S3:**
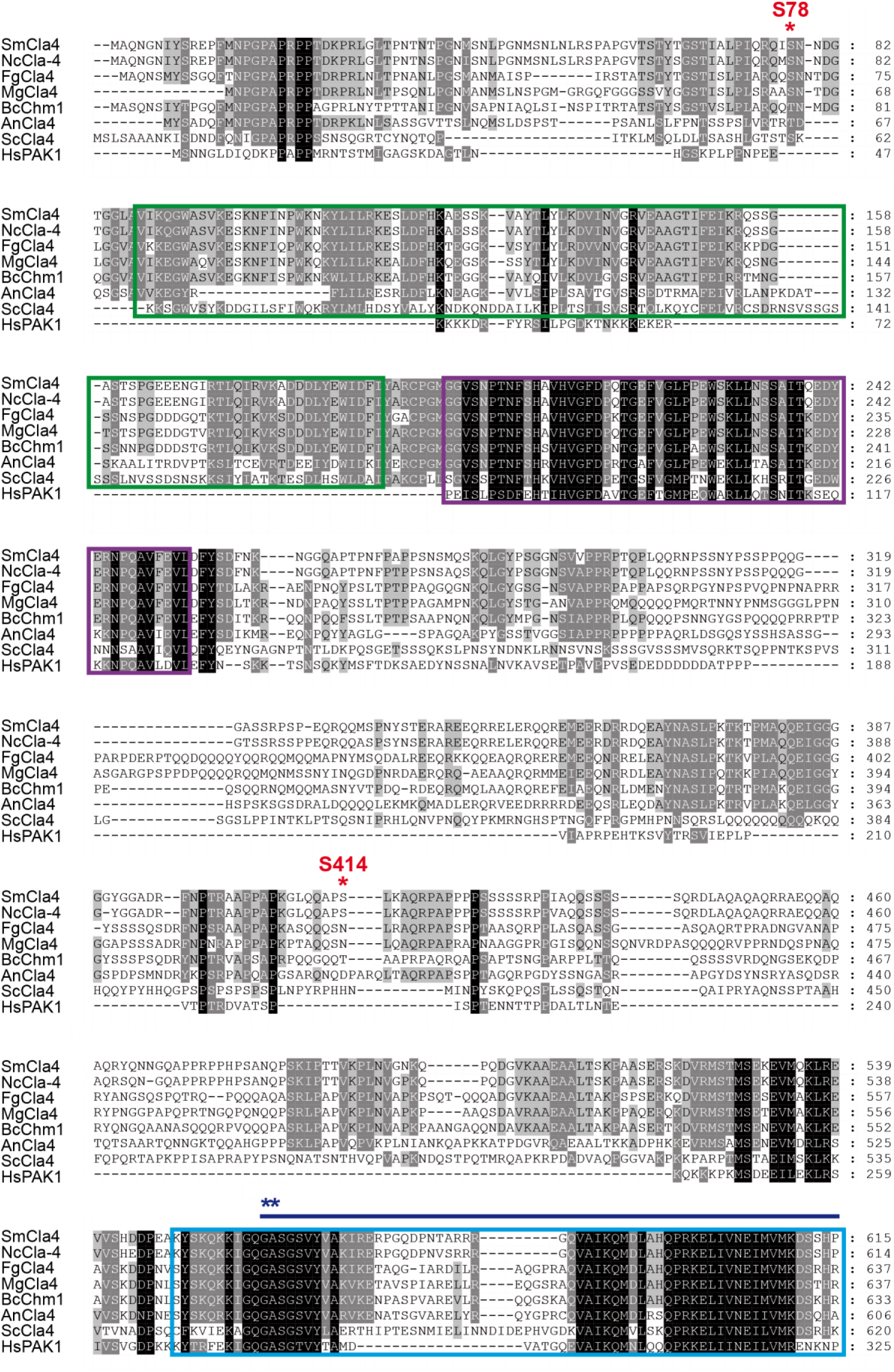
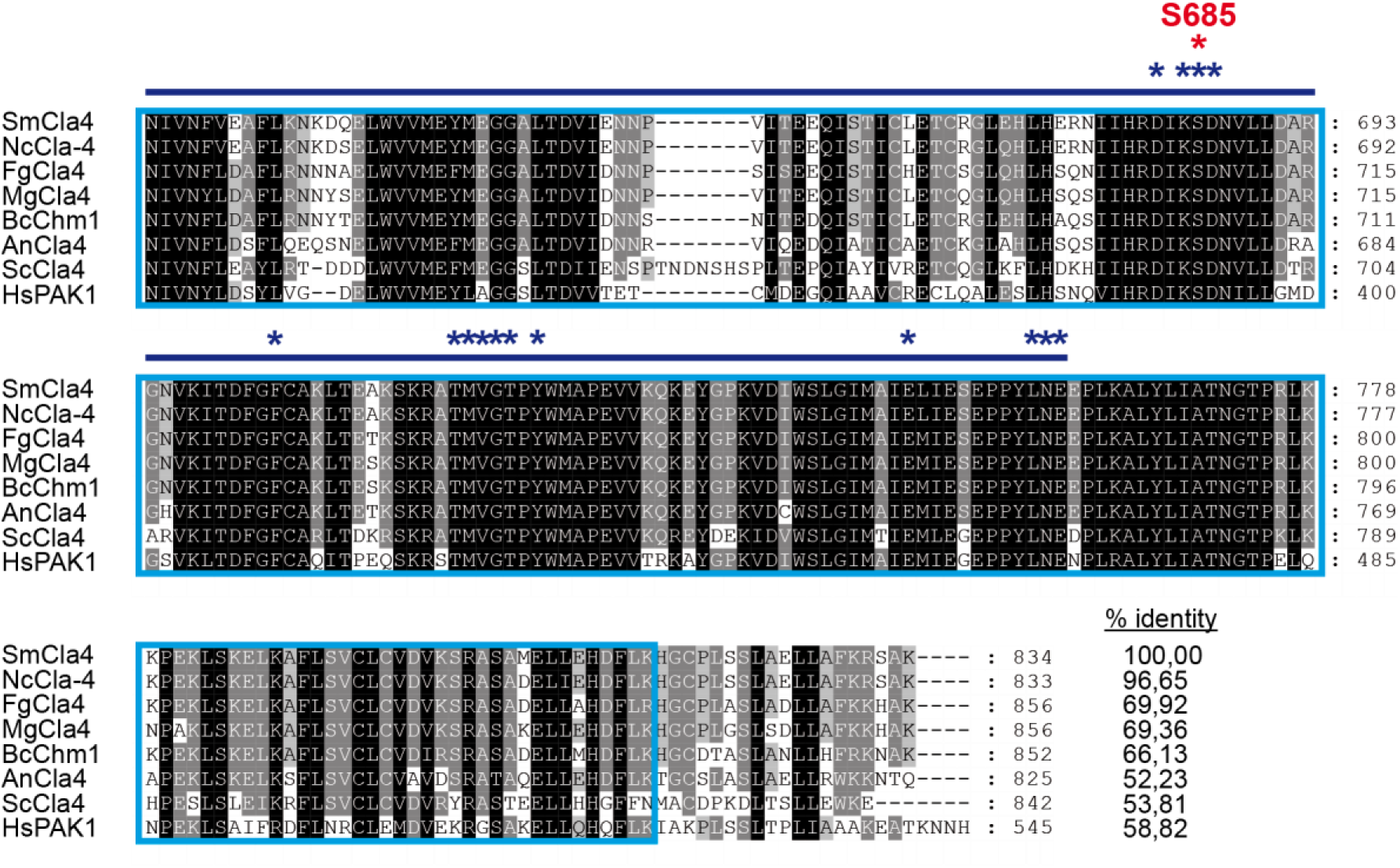
Multiple sequence alignment of CLA4 proteins from filamentous fungi, yeast and human using ClustalW. CLA4 sequences are from the following organisms: *S. macrospora* (Sm, KAA8624319.1), *N. crassa* (Nc, EAA28056.1), *F. graminaerum* (Fg, SCB65154.1), *M. grisea* (Mg, AAL15449.2), *B. cinerea* (Bc, XP_024547628.1), *A. nidulans* (An, EAA60124.1) as well as *S. cerevisiae* (Sc, CAA96216.1) and *H. sapiens* (Hs, AAC50590.1). The p21-binding domain (PBD) (199-253) and the kinase domain (549-815) are displayed by purple and light blue boxes, respectively. The pleckstrin homology (PH) domain (88-190) is shown by a green box. Red and dark blue asterisks indicate the identified phosphorylation sites of *S. macrospora* CLA4 and the substrate binding sites, respectively. % identity shows the percentage of identical amino acids compared to *S. macrospora* CLA4.

**Figure S4:**
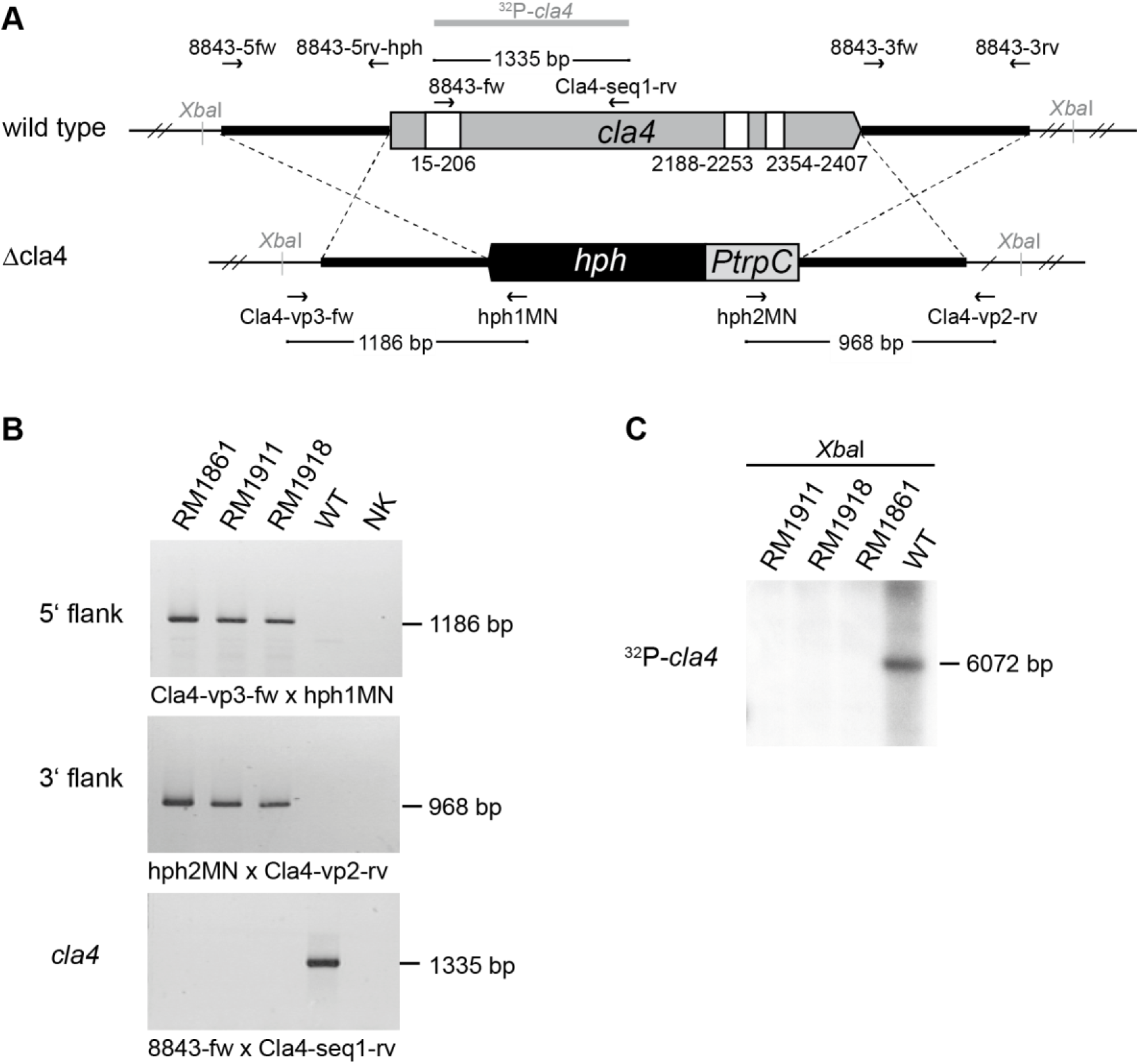
Generation and verification of Δcla4 deletion strains. (A) Genomic region of the *cla4* locus in the wild type and the Δcla4 deletion strain. Arrows mark oligonucleotides used for construction of the deletion plasmid and for verification of the deletion. Black lines indicate PCR fragments, generated with oligonucleotides as shown. Sites of restriction enzyme and the probe used for Southern Blot analysis are indicated by grey letters and a bold grey line, respectively. (B) PCR analyses for verifications of the Δcla4 deletion strains RM1861, RM1911 and RM1918. The homologous integration of the 5’ and 3’ flank and the *cla4* deletion was verified by PCR using the indicated oligonucleotides. Wild type served as a control, and the negative control (NK) contained no genomic DNA. (C) *Xba*I restricted genomic DNA served for Southern hybridization analysis with the radioactively labelled *cla4* specific probe. Genomic DNA was obtained from RM1861, RM1911 and RM1918 as well as the wild type as control. Abbreviations: *hph*, hygromycin B resistance gene from *E. coli*; *PtrpC*, constitutive *trpC* promotor from *Aspergillus nidulans*.

**Figure S5:**
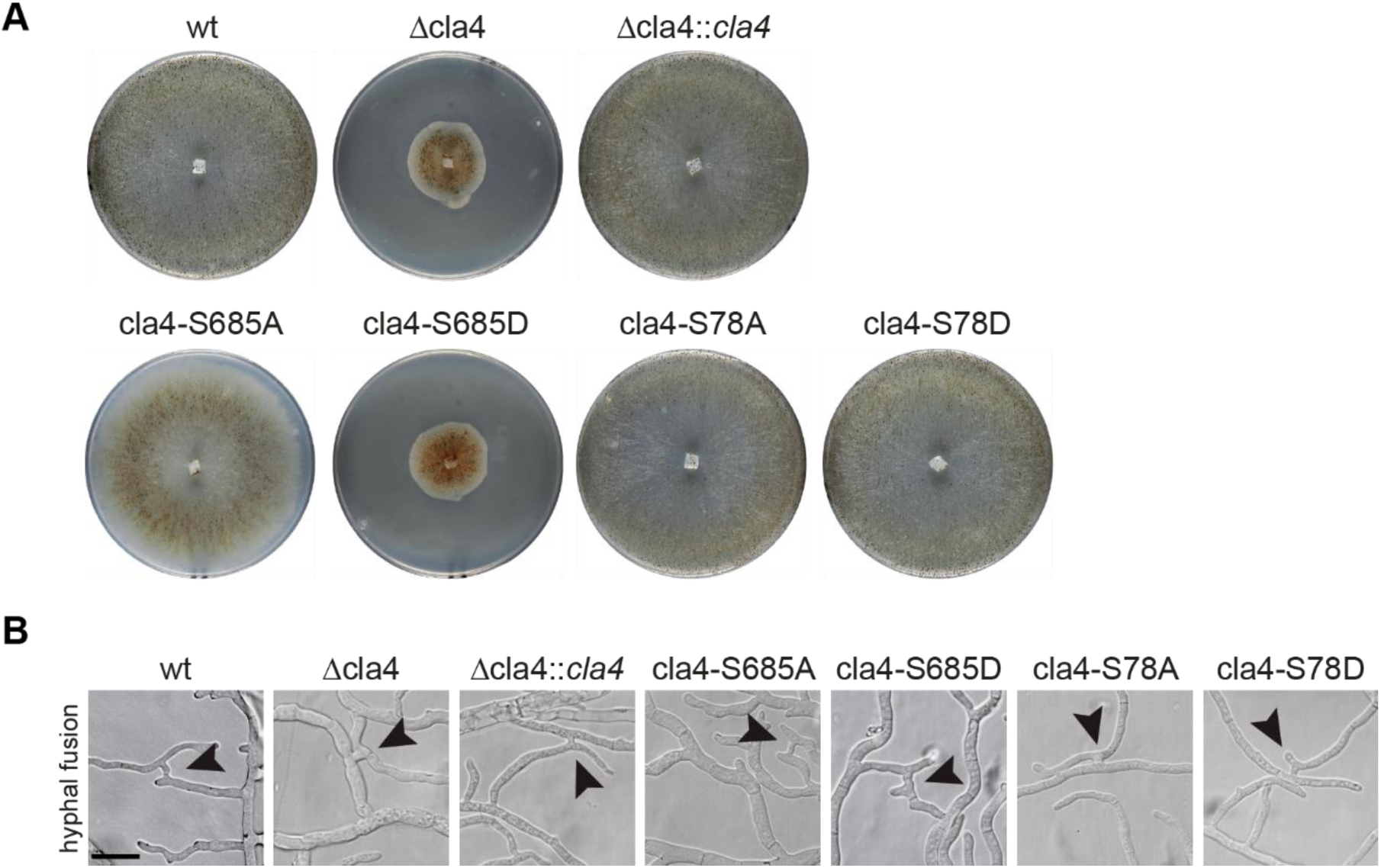
Analysis of vegetative growth and hyphal fusion. As indicated, images were obtained from wild type, Δcla4, and transformants, carrying wild type, phospho-mimetic or phospho-deficient versions of *cla4*. All gene constructs were transferred into Δcla4 strain. (A) Documentation of the colony morphology of strains grown on BMM medium at 27°C for seven days. (B) Investigation of hyphal fusion events in a region 5-10 mm of the colony edges. Arrowheads mark hyphal anastomosis. Strains were grown on cellophane-coated MMS medium at 27°C for 2-4 d. Scale bar is 20 μm. In all cases, we show a single representative strain out of at least three individual transformants.

